# Linking Early Bacterial Responses to Growth Inhibition via Glucose-Turbidity Dynamics for Rapid Antimicrobial Susceptibility Testing

**DOI:** 10.64898/2026.11.19.767763

**Authors:** Yuanzhuo Deng, Shaohan Xu, Huan Hu, Haitao Yang, Wenqian Jiang, Tianzhu Zhu, Qing Li, Jun Lin

## Abstract

Earlier susceptibility information can guide timely antibiotic selection, while metabolic measurements can broaden the characterization of bacterial drug responses. We developed glucose-turbidity antimicrobial susceptibility testing (GT-AST) to pair residual-glucose assay signals with turbidity and relate early responses to growth inhibition. Among 43 cultured isolates, some antibiotic-exposed cultures showed glucose-associated changes despite attenuated turbidity responses. These observations provide physiological clues for studying metabolic activity when growth is limited, including future investigations of viable but non-culturable states or persister cells. Conventional broth microdilution relies on overnight growth readings; GT-AST used trajectories collected during approximately 2 h of antibiotic exposure to predict subsequent growth and estimate minimum inhibitory concentrations (MICs). Across 100 repeats of isolate-grouped cross-validation, essential agreement between predicted MICs and independently measured reference MICs averaged 87.1%. Eight bronchoalveolar lavage fluid specimens were tested directly without prior bacterial isolation. Levofloxacin susceptibility classifications based on observed growth agreed with hospital results in all four culture-positive specimens. Exact and essential agreement between predicted and hospital MICs each averaged 99.0%. Together, these findings support developing GT-AST to provide earlier susceptibility information for effective, appropriately targeted treatment of patients.

## 1. Introduction

For patients with bacterial infections, timely selection of effective and appropriately targeted antibiotics requires early access to reliable information about the drug responses of the infecting bacteria. Among 2,418 intensive care patients with hospital-acquired bloodstream infection, 28-day mortality was 32.8% with and 40.0% without appropriate antimicrobial therapy within 24 h after blood culture collection (Loiodice et al., 2024). Phenotypic antimicrobial susceptibility testing (AST) informs antibiotic selection by measuring drug-induced growth inhibition. The minimum inhibitory concentration (MIC) is the lowest concentration that inhibits visible growth under defined conditions, and clinical interpretation relates susceptibility to the likelihood of therapeutic success at a specified exposure. Conventional broth microdilution (BMD) commonly requires overnight incubation for rapidly growing bacteria (EUCAST, 2026). Phenotypic measurements during early antibiotic exposure could provide an earlier basis for assessing growth inhibition and guiding treatment selection for patients.

Early measurements can also broaden the physiological characterization of bacterial drug responses. Growth and metabolic activity reflect distinct aspects of bacterial physiology: glucose uptake can continue after growth arrest under non-carbon nutrient limitation (Chubukov and Sauer, 2014), and experiments that uncouple metabolic state from growth have shown that this distinction matters for antibiotic lethality (Lopatkin et al., 2019). These findings provide a rationale for examining glucose utilization alongside growth-associated turbidity changes during antibiotic exposure. Such paired observations could reveal metabolic responses accompanying limited population growth and provide physiological clues for future studies of persister and viable but non-culturable (VBNC) populations (Balaban et al., 2019; Bamford et al., 2017).

Rapid phenotypic AST studies have shown that early growth and other physiological responses can inform susceptibility assessment (Baltekin et al., 2017; Zhang et al., 2020; Sturm et al., 2024). In particular, glucose consumption has been monitored by infrared spectroscopy and included among species-dependent metabolic markers measured by mass spectrometry (Jafari et al., 2024; Rydzak et al., 2022). These precedents motivate a paired comparison of glucose and turbidity responses within the same antibiotic exposure: whether reduced turbidity change is accompanied by reduced glucose utilization, and whether their early trajectories can predict subsequent growth inhibition across antibiotic concentrations and support MIC estimation.

Measuring early drug responses directly in clinical specimens could reduce the waiting period associated with prior isolation, allowing susceptibility information to inform treatment selection sooner. Direct gradient diffusion testing of bronchial aspirates and automated microscopy of mini-bronchoalveolar lavage (mini-BAL) specimens have explored this route to susceptibility assessment (Burillo et al., 2019; Douglas et al., 2015). These precedents motivate evaluating whether paired glucose and turbidity responses can be measured directly in bronchoalveolar lavage fluid (BALF). Comparing susceptibility classifications from direct specimen testing with hospital results could then help assess the potential of this approach to inform timely antibiotic selection.

Here, we present glucose-turbidity antimicrobial susceptibility testing (GT-AST), which pairs residual-glucose assay signals with turbidity during approximately 2 h of antibiotic exposure to relate early drug responses to growth inhibition. Using 43 cultured isolates, we characterized the relationship between the two responses across antibiotic concentrations and evaluated early trajectories for predicting growth assessed manually after overnight incubation. MICs estimated from these predictions were compared with independently measured reference MICs. Anticipating that some BALF specimens would contain sufficient bacteria for testing, we initiated paired measurements in eight specimens without prior bacterial isolation and retrospectively evaluated growth predictions. Levofloxacin susceptibility classifications based on observed growth showed complete agreement with hospital results for all four culture-positive specimens. We also compared MICs estimated from the BALF growth predictions with hospital MICs to examine whether early responses could support MIC assessment directly from patient specimens.

## 2. Materials and methods

### 2.1 Bacterial isolates and study material

The cultured-isolate collection comprised 43 isolates: 8 *Escherichia coli*, 12 *Klebsiella pneumoniae*, 6 *Acinetobacter baumannii*, 6 *Pseudomonas aeruginosa* and 11 *Staphylococcus aureus*. These included 41 clinical isolates and two reference strains, *E. coli* ATCC 25922 and *S. aureus* ATCC 29213. Species identification used a Bruker Microflex LT mass spectrometry system, with 16S rRNA gene sequencing for uncertain identifications.

The cultured-isolate experiments yielded 1,605 paired observations: 1,400 antibiotic-exposed observations and 205 no-drug controls. Each paired observation comprised the glucose and turbidity time courses for one isolate at a given drug concentration or under the no-drug condition.

Eight BALF specimens from six patients in hospital intensive care units (ICUs) and general pulmonary wards were tested directly. The BALF specimens and the clinical isolates came from separate patient groups.

### 2.2 Pure-culture preparation and antibiotic exposure

Clinical isolates were purified on Columbia blood agar and stored at −80 °C. Before testing, isolates were recovered on blood agar at 37 °C for 12-16 h. Colonies were suspended in 0.45% NaCl and adjusted to 0.5 McFarland. The standardized suspension was diluted tenfold to prepare the working suspension. Antibiotic exposure was performed in Mueller-Hinton (MH) broth at a final glucose concentration of 3 g/L. The working suspension was mixed with this broth containing the test antibiotic at a suspension-to-broth ratio of 1:19 (v/v), giving a target final inoculum of approximately 5 × 10□ colony-forming units (CFU)/mL. After mixing, cultures were distributed into two 96-well deep-well plates. Plates were centrifuged at 1,000 × g for 30 s and incubated at 37 °C with shaking at 700 rpm. No-drug growth controls and uninoculated controls were included.

Across the pure-culture experiments, single-agent concentrations followed twofold series over 2-32 µg/mL for amikacin; 1-64 µg/mL for aztreonam, cefazolin and ceftriaxone; 0.25-4 µg/mL for ciprofloxacin; 0.25-8 µg/mL for erythromycin; 0.5-8 µg/mL for levofloxacin (LEV); and 2-256 µg/mL for nitrofurantoin. Ampicillin-sulbactam was tested in a twofold series of paired final concentrations from 1/0.5 to 64/32 µg/mL; amikacin-levofloxacin was tested at 2/1, 8/4 and 64/32 µg/mL. In each pair, the first and second values denote the final concentrations of the first and second named components, respectively; both combinations used a fixed mass concentration ratio of 2:1.

### 2.3 Paired glucose and turbidity measurements

Eight samples were collected at 15-min intervals from 30 to 135 min after inoculation of the antibiotic-containing culture system, with four samples taken from each of the two deep-well plates. The first reading, obtained at 30 min, was denoted T1. At each time point, a 200 µL culture aliquot was transferred to a fresh 96-well plate and read at 620 nm using a microplate reader to measure turbidity. The aliquot was then centrifuged at 3,000 × g for 2 min, and the supernatant was used for glucose measurement by o-toluidine colorimetry (Hultman, 1959). A 40 µL volume of supernatant was mixed with 150 µL 6% o-toluidine reagent in a 96-well PCR plate, heated at 95 °C for 12 min and centrifuged at 3,000 × g for 2 min. A 150 µL aliquot of the resulting supernatant was transferred to a plate for absorbance measurement at 620 nm. Both trajectories were analyzed in absorbance units.

A single calibration curve was fitted to glucose standards in MH medium over 0-440 mg/dL. Technical repeat wells from the color-development experiments were used to assess assay precision.

### 2.4 Manual assessment of growth

Growth was assessed manually after overnight incubation. Wells with visible turbidity were labeled 1 (growth), and wells without visible turbidity were labeled 0 (no growth). These labels served as the reference outcome for growth prediction at each tested concentration.

### 2.5 Direct testing of BALF

The eight BALF specimens entered the assay directly, without prior isolation. BALF was mixed with glucose-supplemented MH broth at 1:9 (v/v) and distributed across two 96-well deep-well plates at 1 mL per well. Cultures were incubated at 37 °C with shaking at 700 rpm. Levofloxacin concentrations were 0, 0.5, 1, 2, 4, 6 and 8 µg/mL, with three technical replicates per concentration, yielding 168 paired observations: 144 antibiotic-exposed observations and 24 no-drug controls. Sampling followed the eight time points described above, with four 200 µL aliquots taken from each well on each plate. The remaining 200 µL per well was incubated overnight for manual growth assessment.

In parallel with GT-AST, separate aliquots of the original BALF specimens were serially diluted, spread on sheep blood agar at 100 µL per plate and incubated overnight at 37 °C to examine the inoculum background. The tenfold assay dilution was selected in separate preliminary plating experiments, using the cultured-isolate inoculation conditions as a reference. Inoculum adequacy was assessed using the plating results together with each specimen’s no-drug growth outcome. Species identities used for the full model were obtained from subsequently cultured isolates.

### 2.6 Reference MIC measurements

For cultured isolates, reference MICs were obtained independently of GT-AST from hospital clinical laboratories using an automated AST system (VITEK 2, BioMérieux). Selected results showing dilution-step discrepancies were rechecked by broth dilution and E-test. For amikacin-levofloxacin (AMK_LEV), reference MICs were obtained by independent retesting of the drug combination using a broth method.

Reference MICs were available for all 228 isolate-drug combinations.

### 2.7 Resource accounting

Reagent costs were estimated from purchase prices and estimated quantities for one drug concentration series sampled at eight time points per BALF specimen (Supplementary Table S6).

### 2.8 Control-referenced response analysis

The paired trajectories were used to characterize drug responses and to predict subsequent growth. Cultured-isolate observations supported response analysis and model development, with manually assessed growth as the prediction target. The resulting models were applied retrospectively to BALF specimens, and MIC estimates derived from cultured-isolate and BALF predictions were compared with hospital MICs.

Glucose decreases and turbidity increases were defined from the means of the initial and terminal reading windows using Eqs. (1) and (2):

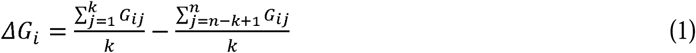

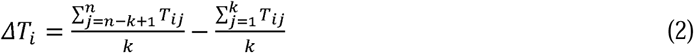

Here i = 1, …, N indexes the N paired observations, each representing one well condition; j = 1, …, n indexes the ordered sampling points. G□□ and T□□ are the glucose and turbidity readings for observation i at point j, respectively. ΔG□ and ΔT□ denote the glucose decrease and turbidity increase. The positive integer k is the number of readings averaged at each end (1 ≤ k ≤ n). Readings and changes are expressed in absorbance units. This study used n = 8 and k = 2, so the terminal window comprised sampling points 7 and 8.

No-drug reference pools were formed by species and antibiotic. Reference changes were the median ΔG□ and ΔT□ within the corresponding pool, with denominator floors of 0.01 and 0.005 absorbance units, respectively. Each response score was calculated as the corresponding change divided by twice its reference change and clipped to the interval 0-1.2. Individual exposed trajectories were also compared with the matched no-drug well from the same isolate and experimental plate.

Response screening included single-agent exposed observations supported by at least five no-drug reference observations and without a signal-quality flag. A glucose flag was assigned when an internal reading differed from the mean of its two neighbors by more than 0.05 absorbance units, or when two successive readings beginning at an internal time point were below 0.06 and the immediately preceding reading was above 0.10. Turbidity flags identified an initial reading exceeding both the second reading and the median of subsequent readings by more than 0.015, or an internal peak exceeding each neighbor by more than 0.03. Within this subset, glucose scores ≥0.5 together with turbidity scores ≤0.3 served as exploratory screening thresholds for relatively preserved glucose responses with attenuated turbidity responses. Manual review was used to choose the illustrative cases. For each selected isolate-drug series, all eligible measured concentrations were retained, with manually assessed growth recorded separately.

### 2.9 Ordination of paired trajectories

Principal coordinates analysis (PCoA) included the observations in the response map and quality-eligible combination-drug observations. Each observation was represented by 22 features summarizing within-channel changes, ranges and slopes of the paired trajectories. For the series-level analysis, these features were summarized across tested concentrations within each isolate and complete drug context, with the number of concentrations appended, giving 89 features per series.

Features were standardized separately at the observation and series levels using the population mean and standard deviation. Euclidean distances were calculated from the complete standardized feature matrix, and classical PCoA was performed by eigendecomposition of the double-centered squared-distance matrix (Gower, 1966). The first two positive axes were displayed. Display groups distinguished selected observations from the remaining observations and series containing at least one selected observation from the remaining series. The additional combination-drug observations supplied background trajectories. Ordination used features derived from the paired trajectories and, at the series level, the number of measured concentrations. Group R² was calculated as the between-group fraction of the total distance-based sum of squares (Anderson, 2001). Ordination and these group summaries were used descriptively for signal-derived groups with repeated observations from the same isolates.

### 2.10 Model inputs and architecture

The full GT-AST model received five input blocks: species indicators, antibiotic-component indicators, concentration-level indicators, glucose trajectories and turbidity trajectories. The indicator blocks contained 11, 10 and 10 entries, respectively. Species and concentration levels were one-hot encoded; combination treatments activated both relevant antibiotic-component indicators. Concentration level 0 denoted no drug; levels 1-9 were categorical identifiers ordered within each complete drug context, with the same level potentially representing different actual concentrations across contexts (Supplementary Table S9). MIC derivation and dilution-step comparisons used actual concentrations. For the full model, each trajectory was centered on its first reading:

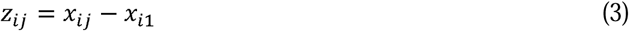

Here x□□ is the reading of one channel for observation i at sampling point j, x□□ is that observation’s first reading of the same channel, and z□□ is the centered reading. The indices i = 1, …, N and j = 1, …, n retain the preceding definitions. Centering was applied separately to glucose and turbidity, retaining absorbance units. This study used n = 8 and retained all eight sampling points. Each resulting time column was then standardized using only the observations assigned to model optimization. Means and population standard deviations were estimated separately for each channel and time column; a standard deviation below 10□□ was replaced by 1. The same fitted transformation was applied to the corresponding stopping, held-out and BALF observations. Indicator blocks were left unscaled.

Glucose and turbidity trajectories were encoded by separate one-dimensional convolutional networks, while each indicator block used a fully connected encoder. The five encoded vectors were combined by two-head attention and passed to a fully connected prediction head. A sigmoid transformation of its output logit gave the model growth score.

### 2.11 Training and isolate-grouped evaluation

Models were trained using AdamW (Loshchilov and Hutter, 2019) and binary cross-entropy loss, with the positive class weighted by the ratio of negative to positive observations in the optimization partition. Each model used a seven-fold StratifiedGroupKFold split with isolate grouping for optimization and early stopping; the first split containing both outcome classes in both subsets was selected. Early stopping monitored area under the receiver operating characteristic curve (ROC-AUC), and the weights at the best stopping epoch were retained.

Development performance was evaluated using 100 repeats of five-fold StratifiedGroupKFold with a new allocation of isolates to folds in each repeat. All observations from an isolate, including different drugs, concentrations and repeated wells, remained in the same outer fold. Within each outer training subset, each input configuration was fitted five times using separate isolate-grouped stopping partitions. The arithmetic mean of the five model outputs provided the growth score for each held-out observation. For BALF evaluation, each input configuration was fitted separately five times in each repeat using isolate-grouped optimization and stopping subsets drawn from the full pure-culture development cohort; the mean of the five outputs provided each well’s growth score in the eight specimens. The BALF specimens formed a separate cohort for retrospective feasibility evaluation.

For the full model, a growth score ≥0.5 predicted growth, with manually assessed growth as the positive reference class. Performance was summarized using ROC-AUC, average precision (AP), sensitivity, specificity, precision, accuracy, balanced accuracy and confusion-matrix counts:

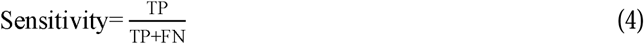

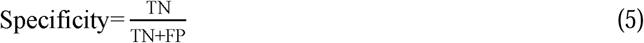

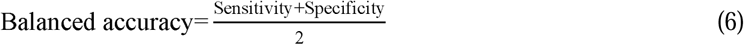

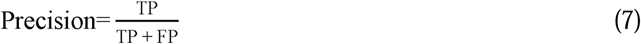

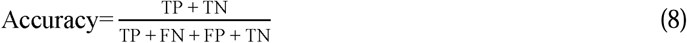

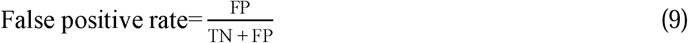

Here true positives (TP) and false negatives (FN) are growing observations predicted as growth and no growth, respectively; true negatives (TN) and false positives (FP) are non-growing observations predicted as no growth and growth, respectively. Sensitivity (recall; Eq. (4)) is the proportion of manually assessed growth observations predicted as growth; specificity (Eq. (5)) is the proportion of manually assessed no-growth observations predicted as no growth. Precision (Eq. (7)) is the proportion of predicted growth observations that were manually assessed as growth. Accuracy (Eq. (8)) is the proportion of observations classified correctly. Balanced accuracy (Eq. (6)) is the arithmetic mean of sensitivity and specificity. These metrics are dimensionless proportions. The ROC curve plots sensitivity against the false-positive rate (Eq. (9)) across score thresholds. Average precision summarizes precision weighted by increases in recall. Development ranking metrics were calculated from pooled out of fold (OOF) scores and also reported separately by outer fold. BALF metrics were calculated from the averaged growth scores. All-observation, antibiotic-exposed and no-drug subsets were reported separately, and individual BALF errors were returned to their specimen and concentration positions.

BALF growth prediction and MIC estimation used 0, 0.5, 1, 2, 4 and 8 µg/mL levofloxacin (144 wells: 120 drug-exposed wells and 24 no-drug controls).

Eight input configurations were fitted with their corresponding encoders using the same isolate partitions, seeds and training settings, and evaluated at a shared threshold of 0.5. These comprised the full model; the full model with the species, glucose, turbidity, antibiotic-component or concentration-level block omitted in turn; the three indicator blocks alone; and the glucose and turbidity trajectories alone.

### 2.12 Comparison of trajectory representations

Four trajectory representations were compared while holding the observations, isolate partitions, indicator blocks, architecture and training settings constant: original readings, T1-centered readings (Eq. (3)), first differences (Eq. (10)) and second differences (Eq. (11)). The two difference orders were defined as:

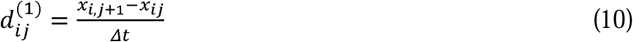

Here d□□□¹□ is the first difference for observation i between sampling points j and j + 1, where i = 1, …, N and j = 1, …, n − 1. The readings x□□ and sampling-point count n retain their preceding meanings. The positive sampling interval Δt is measured in minutes, so d□□□¹□ has units of absorbance per minute.

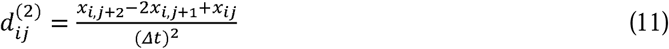

Here d□□□²□ is the second difference for observation i over sampling points j, j + 1 and j + 2, with j = 1, …, n − 2; i, x□□, n and Δt retain the same meanings. Its units are absorbance per minute squared. This study used n = 8 and Δt = 15 min, giving eight original or centered readings, seven first differences and six second differences per channel.

Differences were calculated before training-data standardization. Subtracting the first reading leaves both difference orders unchanged, so only the four distinct representations were fitted in each repeat. Each representation was evaluated using both a common threshold of 0.5 and its own threshold selected to maximize balanced accuracy over pooled development out of fold scores within each repeat; ties were resolved by proximity to 0.5 and then the lower value. The selected thresholds were also applied to the corresponding BALF scores in the same repeat. Both sets of classification results are reported in Supplementary Table S3. The same development-set out of fold predictions were used for both threshold selection and performance evaluation.

### 2.13 Derivation of predicted MICs

For each isolate-drug combination, the full model’s out of fold growth predictions were ordered by increasing antibiotic concentration to derive the predicted MIC. The no-drug control had to be predicted as growing, and replicate predictions had to agree at each concentration. Series failing these conditions were recorded as non-reportable. A skipped concentration was defined by a no-growth prediction at a lower concentration than one predicted to support growth. For series without skips, the MIC was the next tested concentration predicted as no growth above the highest concentration predicted as growth. For series with skips, the highest tested concentration was assigned as the numerical MIC estimate, and the series was flagged in Supplementary Table S4.

In experiments with direct BALF inoculation, growth predictions from the full model were aggregated by majority vote across three replicate wells at each levofloxacin concentration: at least two wells predicting growth defined a growing concentration. When the control majority predicted growth, MICs were derived using the same concentration-order and skipped-concentration rules.

### 2.14 Statistical analysis

MIC agreement was evaluated by comparing predicted and reference MICs. Exact agreement (EX) was defined as a dilution difference of zero, and essential agreement (EA) as agreement within one twofold dilution (ISO, 2021). For pure cultures, dilution differences (d) were calculated as:

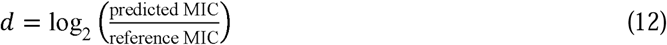

For direct BALF experiments, dilution differences were calculated according to the CLSI Breakpoint Implementation Toolkit (CLSI, 2023). In each evaluation, we calculated classification metrics, derived MICs and assessed MIC agreement for series with a reference MIC. For each metric, we report the mean of the values available across the 100 evaluations. Confidence intervals and paired differences were estimated using 20,000 cluster bootstrap replicates, resampling isolates for pure cultures and patients for BALF, with specimen-level resampling as a sensitivity analysis. Paired comparisons used common bootstrap samples, with models and prediction rules held fixed. The 95% confidence intervals were the 2.5th and 97.5th percentiles of valid replicates. Further details are provided in Supplementary Methods SM5-SM6 and Tables S3 and S8.

The observation was the unit for trajectory and growth-classification summaries, the isolate was the grouping unit for model evaluation and pure-culture MIC resampling, and each isolate-drug series could supply one MIC result per repeat. BALF repeat wells were summarized within their originating specimen, and all specimens from the same patient were resampled together. Modeling used Python 3.12.3 with NumPy 2.3.5, pandas 2.3.3, scikit-learn 1.6.1 (Pedregosa et al., 2011) and PyTorch 2.8.0 (Paszke et al., 2019). Training was performed on the CPU with two PyTorch threads and deterministic algorithms enabled.

## 3. Results

### 3.1 Study design for paired response characterization and growth assessment

Antibiotic selection for patients relies in part on determining whether a drug inhibits bacterial growth. We asked what glucose measurements could add to turbidity-based characterization of bacterial drug responses and how the two early readouts related to subsequent growth. Within a broth-dilution framework, GT-AST paired measurements of residual glucose in culture supernatants with culture turbidity across antibiotic concentrations and a no-drug control during approximately 2 h of early exposure (Fig. 1A,B). The schematic examples illustrate glucose consumption accompanied by increased turbidity or by little turbidity change (Fig. 1C). Each well was assessed manually for visible turbidity after overnight incubation, linking the early response trajectories to a later growth outcome.

**Figure 1.**
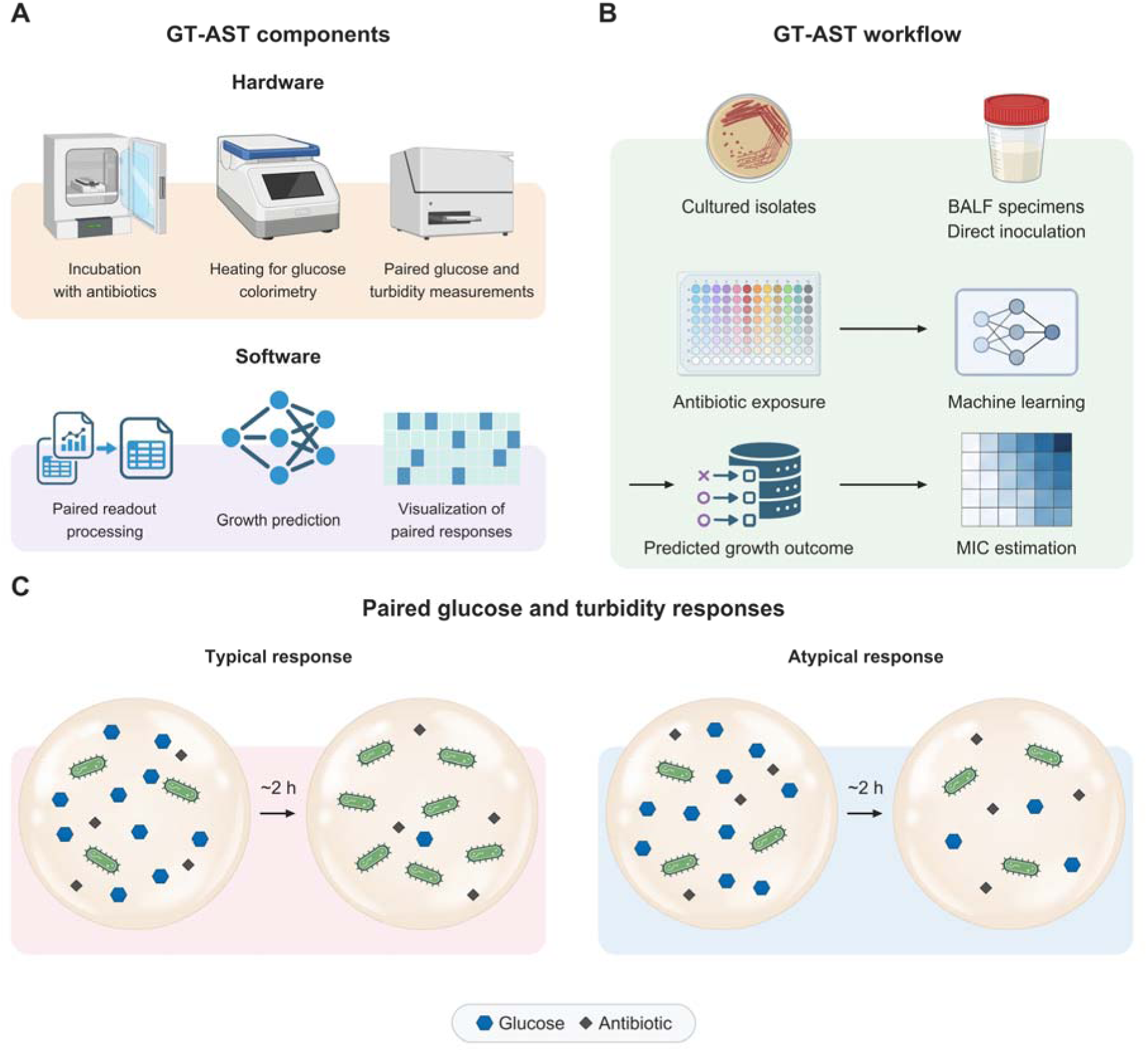
Study design of glucose-turbidity antimicrobial susceptibility testing (GT-AST). A, GT-AST components. Laboratory equipment supports incubation with antibiotics, heating for glucose colorimetry, and paired glucose and turbidity measurements. Software processes the paired readouts, predicts growth, and visualizes paired responses. B, Experimental and analytical workflow. The study used 43 cultured isolates for response characterization and model development, and eight bronchoalveolar lavage fluid (BALF) specimens tested without prior isolation for retrospective feasibility evaluation. Early paired measurements across antibiotic concentrations and a no-drug control provide inputs for growth prediction and minimum inhibitory concentration (MIC) estimation. Growth labels were based on visible turbidity after overnight incubation. C, Typical and atypical paired responses during approximately 2 h of antibiotic exposure, sampled at eight time points from 30 to 135 min after inoculation. In the typical example, glucose consumption accompanies increased turbidity, represented by more bacterial symbols. In the atypical example, glucose consumption accompanies little turbidity change, represented by similar bacterial numbers. Blue hexagons denote glucose and gray diamonds antibiotics.

We used 43 cultured isolates to characterize paired responses and evaluate their use in predicting subsequent growth. The model combined the two trajectories with species, antibiotic and concentration information to predict growth in each well (Fig. 1B), with performance evaluated by isolate-grouped cross-validation. MIC estimates derived from out of fold predictions across eligible concentration series (Fig. 1B) were compared with reference MICs. To explore direct testing of clinical specimens, we also measured paired responses in eight BALF specimens without prior isolation and evaluated their growth predictions retrospectively. This design allowed us to examine the relationship between the two early readouts and then assess their connection to subsequent growth.

### 3.2 Glucose and turbidity capture complementary aspects of antibiotic response

During approximately 2 h of antibiotic exposure, paired measurements revealed differences between the glucose and turbidity responses in two illustrative isolate-antibiotic combinations. In *Klebsiella pneumoniae* study isolate KPN001 exposed to 4 µg/mL ceftriaxone, the glucose assay signal decreased overall, whereas turbidity rose initially and declined after 120 min; turbidity in the matched no-drug well continued to increase (Fig. 2A). In *Acinetobacter baumannii* study isolate ABA001 exposed to 4 µg/mL levofloxacin, the glucose trajectory remained close to that of the matched no-drug control, while turbidity changed little despite increasing control turbidity (Fig. 2B).

**Figure 2.**
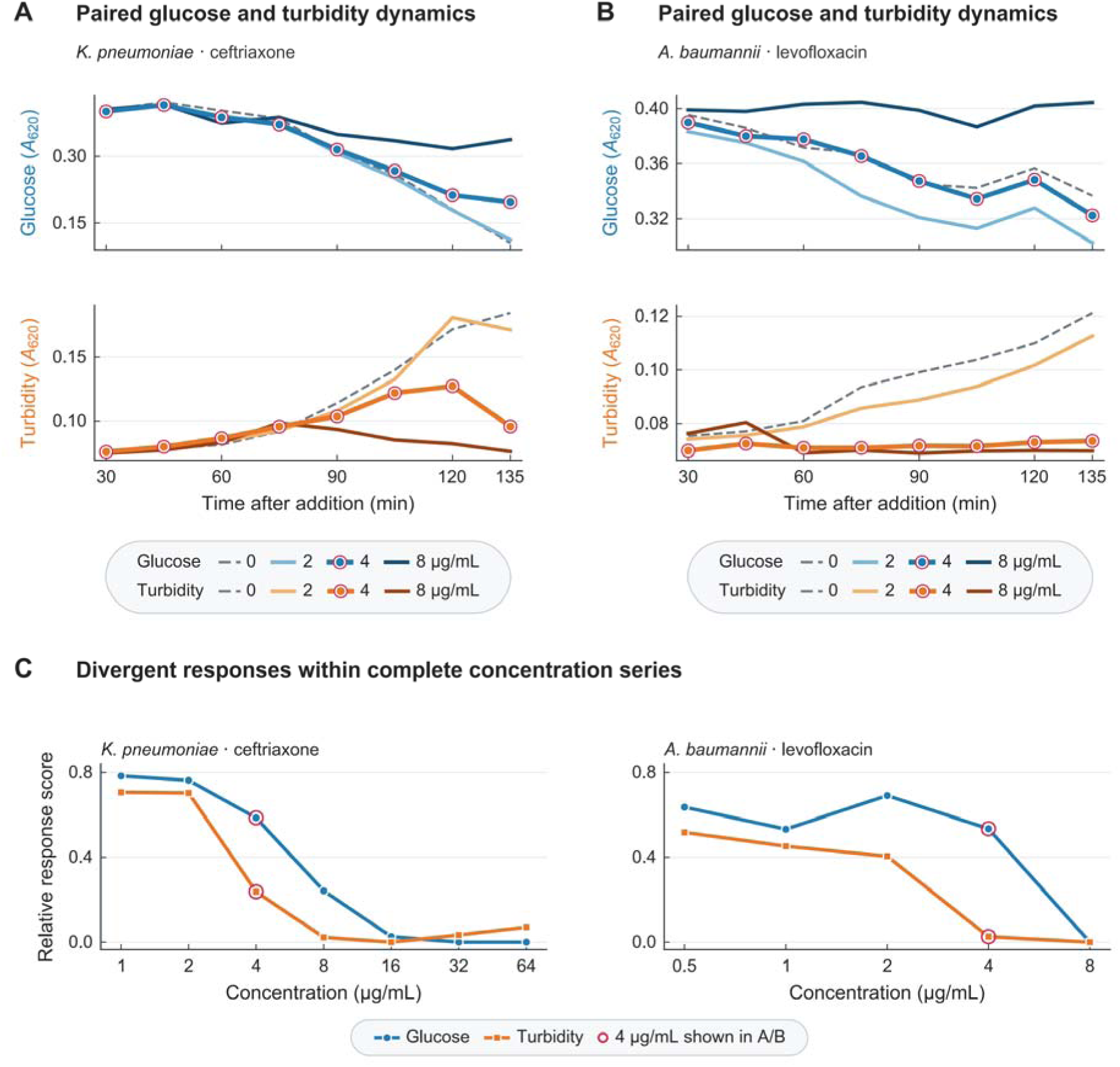
Non-equivalent early glucose and turbidity responses. A,B, Glucose assay signal (upper plots) and culture turbidity (lower plots), both measured as absorbance at 620 nm (A_620_), for *Klebsiella pneumoniae* KPN001 exposed to ceftriaxone (A) and *Acinetobacter baumannii* ABA001 exposed to levofloxacin (B). Each series shows 2, 4 and 8 µg/mL and a matched no-drug control (0 µg/mL), measured at eight time points from 30 to 135 min after inoculation. Each trajectory represents one well. Blue denotes glucose and orange turbidity; darker shades indicate higher antibiotic concentrations. Gray dashed curves show the no-drug well from the same isolate and plate. Thick curves with markers with red outer rings highlight 4 µg/mL, with growth in A and no growth in B. C, Response scores across the complete tested concentration series: 1-64 µg/mL ceftriaxone for KPN001 and 0.5-8 µg/mL levofloxacin for ABA001. Circles denote glucose, squares turbidity, and red rings the 4 µg/mL observations shown in A and B. Scores summarize glucose decreases and turbidity increases relative to no-drug reference pools matched by species and antibiotic: 12 wells from different *K. pneumoniae* isolates for ceftriaxone and six from different *A. baumannii* isolates for levofloxacin. The score calculation, including scaling and clipping, is described in Methods. Growth and no growth were assigned by manual assessment of the presence or absence of visible turbidity after overnight incubation.

Lower absorbance in the colorimetric assay indicates less residual glucose, so a declining signal is consistent with utilization of this carbon source. Continued glucose consumption after growth arrest has been observed in *Escherichia coli* and *Bacillus subtilis* under limitation of nutrients other than carbon (Chubukov and Sauer, 2014). These paired trajectories therefore suggest that glucose-utilizing metabolic activity may persist during early drug exposure even when the turbidity response is attenuated.

To examine how this relationship varied with antibiotic concentration, we compared the complete ceftriaxone series for KPN001 (1-64 µg/mL) and levofloxacin series for ABA001 (0.5-8 µg/mL). Glucose decreases and turbidity increases were summarized as response scores normalized to each readout’s pooled no-drug reference (Fig. 2C). At 4 µg/mL, glucose scores exceeded turbidity scores in both examples. At 8 µg/mL, both scores were lower and their difference was smaller than at 4 µg/mL. In the ABA001 series, increasing levofloxacin from 1 to 2 µg/mL was accompanied by a higher glucose score but a lower turbidity score. These series therefore showed distinct concentration-dependent changes in the glucose-associated and turbidity responses.

To relate these early differences to subsequent growth, we compared the highlighted wells with their manual readings after overnight incubation. For KPN001, the well exposed to 4 µg/mL ceftriaxone became turbid, making this the highest tested concentration with visible growth. For ABA001, the well exposed to 4 µg/mL levofloxacin remained non-turbid, making this the lowest tested concentration without visible growth. Early responses consistent with glucose utilization therefore accompanied both subsequent growth and growth inhibition. The paired measurements complemented the later growth reading by describing the glucose and turbidity responses during drug exposure.

### 3.3 Atypical paired responses accompany different growth outcomes

To assess the distribution and growth outcomes of the response pattern illustrated in Fig. 2, we applied a common exploratory criterion to 985 analyzable drug-exposed observations. Using scores referenced to no-drug controls, the criterion identified relatively preserved glucose responses alongside attenuated turbidity responses (Methods). The observations meeting this criterion (n = 24, 2.4%) included both Fig. 2 examples (Fig. 3A). After overnight incubation, these 24 wells were either turbid (n = 13, 54.2%) or non-turbid (n = 11, 45.8%) (Fig. 3B,D). Thus, this early response pattern occurred in a small subset of the analyzed observations and was followed by either growth or no growth.

**Figure 3.**
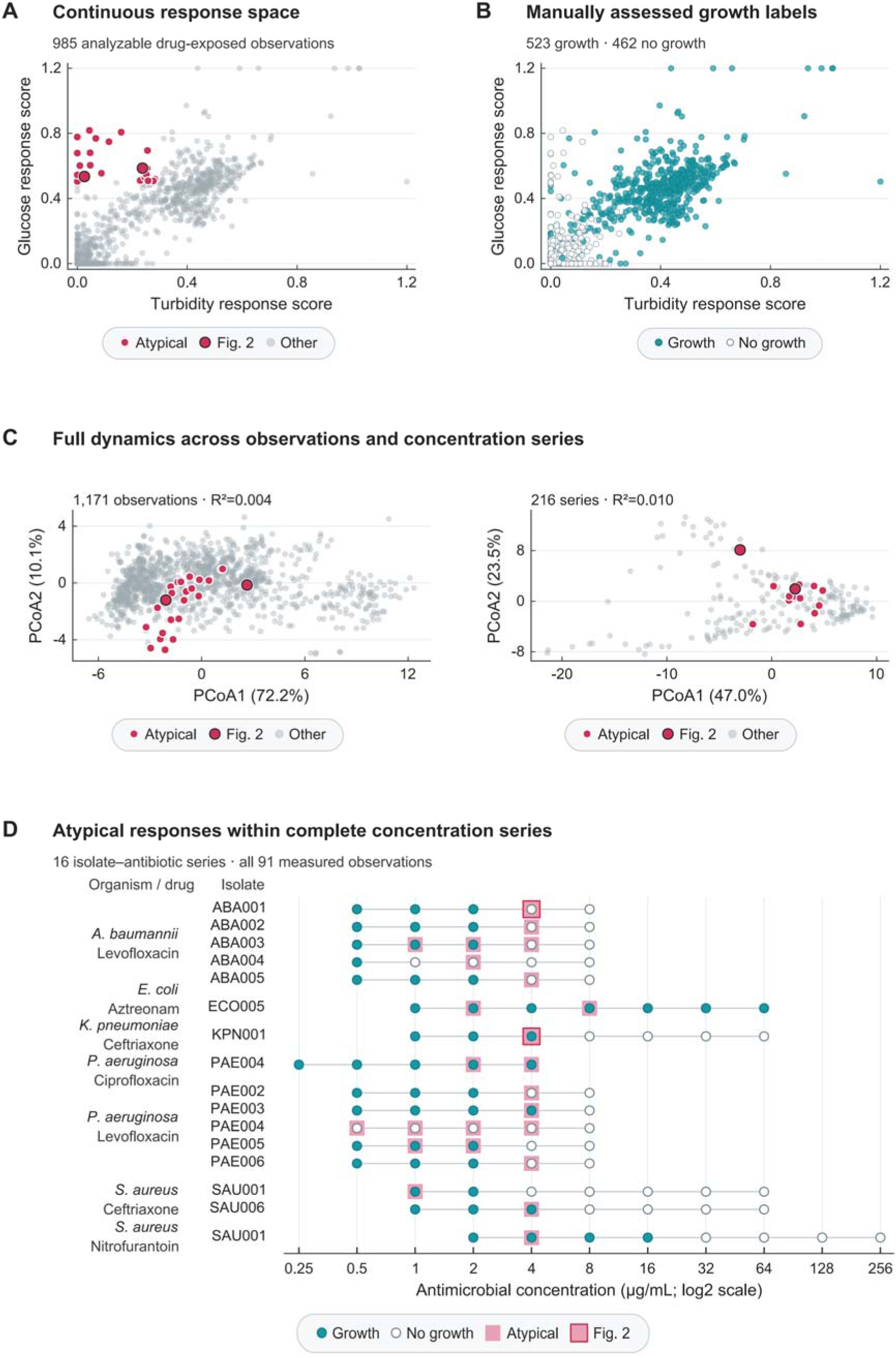
Distribution and growth outcomes of atypical paired responses. A, Control-referenced glucose and turbidity response scores for 985 analyzable drug-exposed observations, each representing one well at one antibiotic concentration. Red points mark the atypical responses (n = 24, 2.4%), operationally defined by glucose scores ≥0.5 and turbidity scores ≤0.3. Larger red points with dark outlines and white halos mark the two Fig. 2 examples within this subset. Score calculation is described in Methods. B, The same observations marked by growth outcome: growth (n = 523, 53.1%) as cyan filled circles and no growth (n = 462, 46.9%) as open circles with gray outlines. C, Exploratory PCoA at the observation level (left; 1,171 observations) and series level (right; 216 isolate-antibiotic series). The analysis includes 186 additional combination-antibiotic observations beyond A and B; single drugs and drug combinations are treated as distinct contexts. Red marks the 24 selected observations or the 16 series containing them; larger red points with dark outlines and white halos mark the Fig. 2 examples or their series. Euclidean distances were calculated from 22 standardized features per observation and 89 per series. The first two axes explain 72.2% and 10.1% of observation-level variation, and 47.0% and 23.5% of series-level variation. Descriptive R² values for selection membership were 0.004 for observations and 0.010 for series. Groups were defined from the same signals, with multiple observations from the same isolates. D, Growth readings across all 91 wells measured across the 16 series containing atypical responses. Cyan filled circles denote growth (n = 50, 54.9%), and open circles with gray outlines denote no growth (n = 41, 45.1%). Pink squares identify the 24 atypical observations with growth (n = 13, 54.2%) or no growth (n = 11, 45.8%). Red square outlines identify the two Fig. 2 examples. Blank positions indicate unmeasured concentrations. Growth and no growth denote the presence and absence of visible turbidity, respectively, on manual assessment after overnight incubation. PCoA, principal coordinates analysis.

To place these contrasting outcomes in their concentration context, we examined all 91 wells measured across the 16 isolate-antibiotic series containing atypical responses (Fig. 3D). These series came from 14 isolates spanning five species and five antibiotics. The observed patterns were a transition from growth to no growth with increasing concentration (n = 13, 81.3%), growth at every tested concentration (n = 2, 12.5%), and no growth at any tested concentration (n = 1, 6.3%). In *Pseudomonas aeruginosa* study isolate PAE004, growth was observed at all five ciprofloxacin concentrations (0.25-4 µg/mL) and at none of the five levofloxacin concentrations (0.5-8 µg/mL). Atypical responses occurred at two ciprofloxacin concentrations (40%) and four levofloxacin concentrations (80%). Thus, the pattern occurred both in series with a growth transition and in series with uniform growth or no growth, including opposite outcomes for the same isolate under different antibiotics.

We next examined whether the selected responses shared a distinct overall dynamic profile. Exploratory PCoA included 1,171 observations and 216 isolate-antibiotic series, with 186 combination-antibiotic observations beyond the screening set (Fig. 3C). The 24 selected observations and the 16 series containing them overlapped with the remaining observations and series on the first two axes. Group membership accounted for approximately 0.4% and 1.0% of the variation in the full observation-level and series-level distance matrices, respectively. One possible explanation is that the control-referenced scores summarize the relative changes in the two readouts, whereas the full trajectory features also capture their amplitudes and time courses. At the series level, a pattern restricted to some concentrations may contribute little to the overall profile. The selected responses accompanied both growth outcomes and overlapped with other response profiles in the displayed dynamic space.

### 3.4 GT-AST uses paired early responses to predict growth and estimate MICs

Because the atypical paired responses in Fig. 3 accompanied both growth and no growth, we asked whether the full trajectories and their experimental context could predict subsequent growth at each drug concentration. The full GT-AST model combined eight glucose and eight turbidity readings from approximately 2 h of antibiotic exposure with species, drug and concentration information. Each trajectory was centered on its first reading, and the prediction target was growth assessed manually after overnight incubation. Development included 1,605 observations from 43 cultured isolates. Five-fold cross-validation kept each isolate’s observations together, so predictions were evaluated on isolates excluded from the corresponding model training.

Across 100 repeats, the pooled out of fold scores distinguished growth from no growth with a mean ROC-AUC of 0.9968 and average precision of 0.9979 (Fig. 4A). Discrimination remained high after excluding no-drug controls: among 1,400 drug-exposed observations, mean ROC-AUC was 0.9960 and average precision was 0.9967. For concentration-series interpretation, scores ≥0.5 were classified as growth. Results across repeats and classification percentages are provided in Supplementary Figs. S4 and S5.

**Figure 4.**
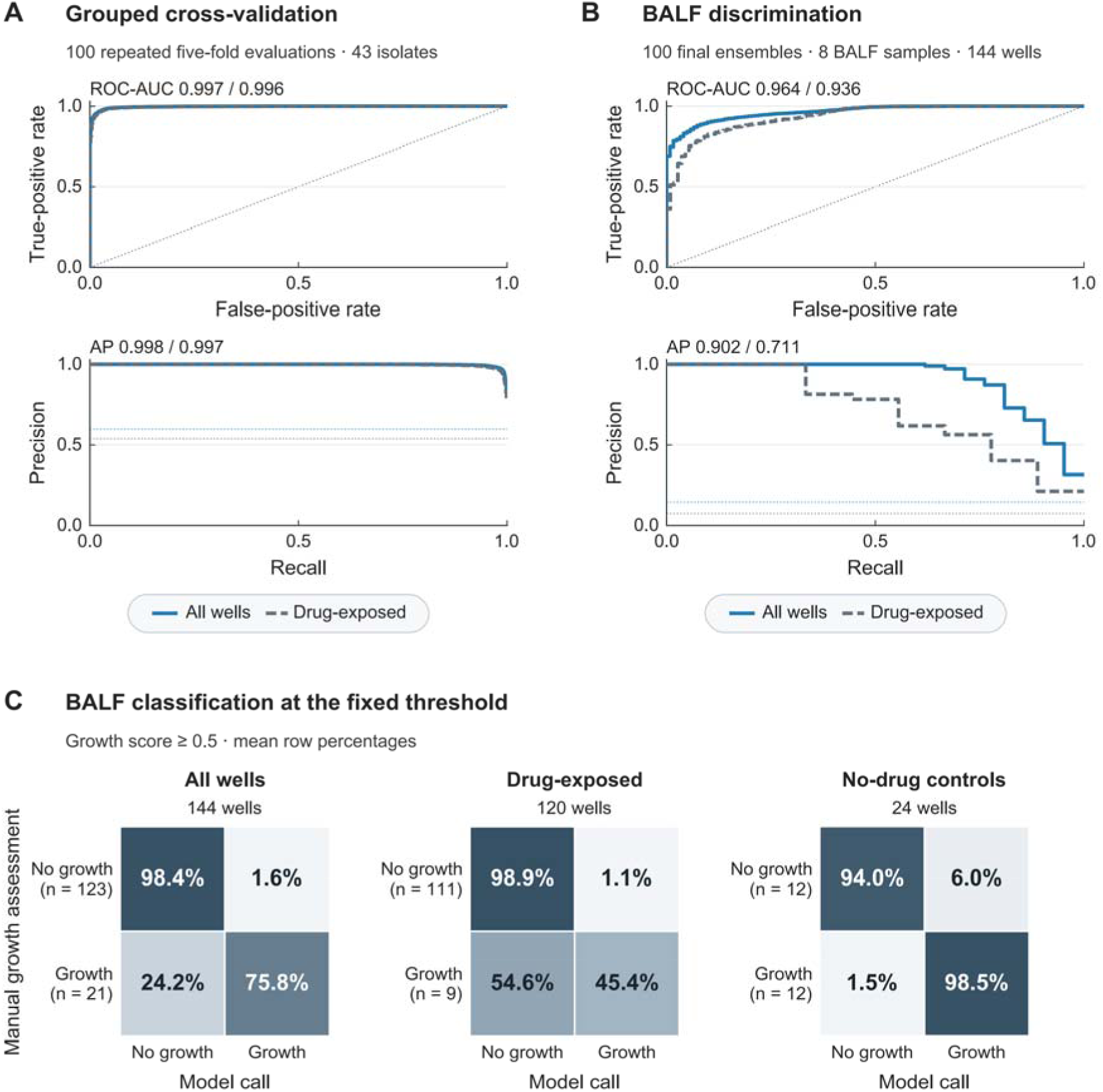
Growth prediction in cultured isolates and directly assayed BALF. Eight glucose and eight turbidity readings at 30-135 min after inoculation are combined with species, drug and concentration information. Each trajectory is centered by subtracting its first reading (T1, 30 min). Development comprises 1,605 observations from 43 cultured isolates. Isolates remain grouped in both outer folds and internal early-stopping partitions. Within each repeat, each held-out score is the mean of five model outputs. For BALF, five models are fitted on the pure-culture development data, and their mean output provides a growth score for each of 144 wells from eight directly assayed bronchoalveolar lavage fluid (BALF) specimens, with species inputs from subsequently cultured isolates. The development cohort included five species categories and the BALF cohort seven, six of which were absent from development. A, Pooled out of fold receiver operating characteristic (ROC; upper) and precision-recall (lower) curves. Blue solid lines show all 1,605 observations and gray dashed lines the 1,400 drug-exposed observations. ROC-AUC values are 0.9968 and 0.9960, and average precision (AP) values are 0.9979 and 0.9967, respectively. B, Corresponding curves for all 144 BALF wells (growth, n = 21, 14.6%) and the 120 drug-exposed wells (growth, n = 9, 7.5%), using the same line codes. Curves and metrics in A and B are means across 100 repeats. ROC-AUC values are 0.9644 and 0.9361, and AP values are 0.9024 and 0.7112, respectively. Dotted ROC lines indicate random ranking; dotted precision-recall lines indicate the growth-label proportion in each subset. C, Confusion matrices at a growth-score threshold of 0.5 for all wells, drug-exposed wells and no-drug controls (left to right). Rows show manual reference outcomes and columns model calls; scores ≥0.5 predict growth. Row labels give the observed well counts. Cell values and blue-gray shading show the mean percentages within each row. Growth denotes visible turbidity on manual assessment after overnight incubation. BALF results are retrospective, with wells nested within eight specimens from six patients. Model outputs are uncalibrated growth scores. Additional performance results are provided in Supplementary Figs. S4, S5 and S7-S9 and Supplementary Tables S3 and S8. AUC, area under the ROC curve.

To assess whether concentration-specific growth predictions could support MIC estimation, we derived MICs from the out of fold predictions. The model predicted MICs for all 228 isolate-drug combinations in 99 of the 100 evaluations and for 226 combinations in the remaining evaluation. Mean MIC prediction coverage was 99.99%. Mean exact agreement with the reference MICs was 64.8% and mean EA was 87.1%. Predictions showed skipped concentrations in 8 to 16 of the 228 combinations per repeat (Supplementary Table S4). Confidence intervals are provided in the Supplementary Results. These comparisons link early growth predictions to independently measured growth-inhibition endpoints.

We next examined growth prediction in directly assayed BALF. Eight specimens from six patients were tested without prior isolation, anticipating that some might contain sufficient bacteria to begin drug-response testing directly. We used the full GT-AST model trained on pure-culture development data to retrospectively evaluate 144 BALF wells. ROC-AUC and average precision averaged 0.9644 and 0.9024 across all wells, and 0.9361 and 0.7112 across the 120 drug-exposed wells (Fig. 4B). Among the 120 drug-exposed wells, overnight readings showed growth (n = 9, 7.5%) or no growth (n = 111, 92.5%). At the 0.5 threshold (Fig. 4C), the model detected three to six of the nine growth wells per repeat (mean detection rate, 45.4%) and misclassified zero to three of the 111 no-growth wells (mean false-positive rate, 1.1%). Detailed confidence intervals are provided in Supplementary Table S3.

To examine growth prediction before species identification, we also evaluated the model without species input, which used glucose and turbidity trajectories, drug and concentration inputs. Among drug-exposed BALF wells, mean growth detection was 47.2% without species input and 45.4% with the full model (Supplementary Table S8). These retrospective results support investigating direct BALF workflows in which phenotypic measurement and species identification proceed in parallel.

Growth detection and false-positive calls also differed between trajectory representations (Supplementary Figs. S7-S9; Supplementary Table S3). Together, these results show that early paired responses can predict subsequent growth across antibiotic concentrations and support MIC estimation in cultured isolates. Direct BALF measurements allow the same link to be examined at the specimen level, from early responses through growth inhibition to comparison with hospital MICs.

### 3.5 GT-AST characterizes paired drug responses directly in BALF

Eight BALF specimens from six patients were tested directly by GT-AST at a final tenfold dilution, without prior isolation. Each specimen was evaluated at five levofloxacin concentrations (0.5, 1, 2, 4 and 8 µg/mL), with three wells per concentration and three no-drug controls. Meanwhile, separate aliquots were plated to examine the cultivable inoculum background. Figure 5A shows dilution plates from BALF001 and BALF004, and Fig. 5B shows their paired glucose and turbidity responses during approximately 2 h of drug exposure.

**Figure 5.**
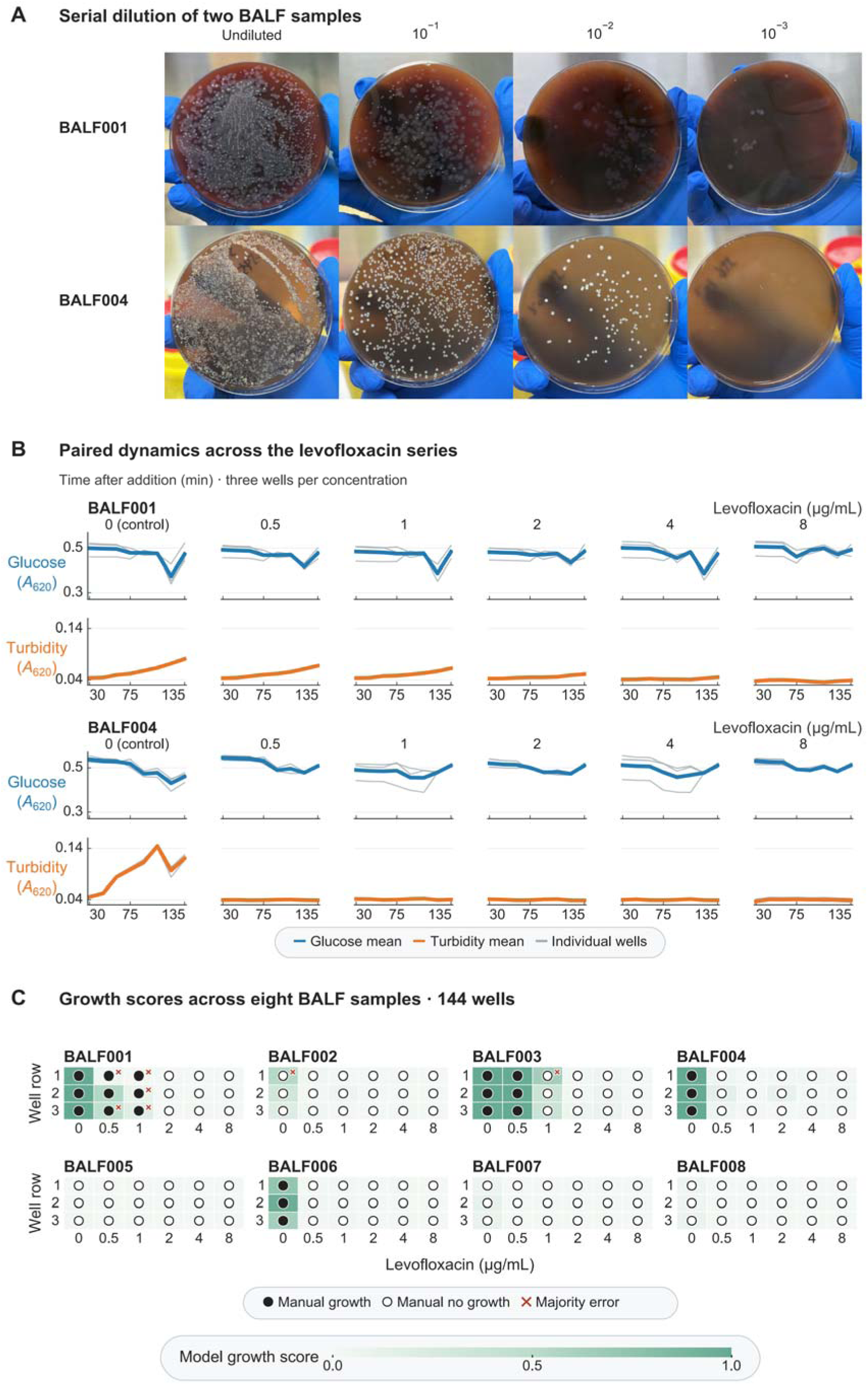
Paired drug responses and growth outcomes in directly assayed BALF. A, Parallel serial-dilution plating of separate aliquots from bronchoalveolar lavage fluid (BALF) specimens BALF001 (upper row) and BALF004 (lower row). Columns show undiluted fluid and 10L¹, 10L² and 10L³ dilutions, with 100 µL plated per plate, to illustrate the cultivable inoculum background. B, Paired glucose assay signal (blue) and turbidity (orange), both measured as absorbance at 620 nm (A_620_), for the two specimens. Glucose-turbidity antimicrobial susceptibility testing (GT-AST) used a final tenfold dilution of the original BALF without prior isolation. Columns show five levofloxacin (LEV) concentrations (0.5, 1, 2, 4 and 8 µg/mL) and the no-drug control (0). Thin gray lines show three technical replicate wells per concentration; thick colored lines show their arithmetic mean. Eight readings were obtained at 30-135 min after inoculation. Lines connect the original measurements without smoothing, with common axes within each readout. C, Model growth scores and manually assessed growth for all 144 wells from eight specimens. The full model uses trajectories centered on each well’s first reading (T1), with species inputs from subsequently cultured isolates. Columns denote 0, 0.5, 1, 2, 4 and 8 µg/mL LEV; rows show the three technical replicate wells at each concentration. Green shading indicates the mean uncalibrated growth score (0-1) across 100 repeats. Solid black circles and white circles with black outlines indicate growth and no growth after overnight incubation, respectively. Scores ≥0.5 predict growth; orange-red crosses mark wells with discrepant calls in more than half of the 100 repeats (n = 7, 4.9% of the 144 wells). Drug-induced inhibition is evaluated in the four culture-positive specimens; the remaining four specimens showed no growth in either control or exposed wells. Alternative representations and calls are shown in Supplementary Figs. S7-S9

The two readouts showed different concentration responses in these specimens. In BALF001, the mean turbidity rise declined with increasing levofloxacin concentration: A_620_ increased from 0.044 to 0.081 without drug and from 0.044 to 0.063 at 1 µg/mL, with little change at 4-8 µg/mL. Glucose signals fluctuated over time across the levofloxacin concentrations. In BALF004, the mean glucose signal without drug fell from 0.536 to 0.460 A_620_ as mean turbidity rose from 0.046 to 0.120. Glucose signals fluctuated in the drug-exposed wells, whereas turbidity changed little at all five drug concentrations (Fig. 5B). Both specimens thus showed glucose-signal variation at concentrations with little early turbidity change.

To relate these early responses to subsequent growth, we examined the concentration series after overnight incubation. Growth was observed in all three no-drug wells in each of BALF001 and BALF004, consistent with their plating results. In BALF001, all three wells became turbid at each of 0.5 and 1 µg/mL. None of the three wells became turbid at any of the higher concentrations (2, 4 and 8 µg/mL). Thus, the attenuated early turbidity rise at 1 µg/mL was followed by visible growth overnight. BALF004 and BALF006 grew only in the no-drug controls; BALF003 grew in the controls and at 0.5 µg/mL. The growing controls in these four specimens provided a within-specimen reference for evaluating drug-induced growth inhibition. In each of BALF002, BALF005, BALF007 and BALF008, all 18 wells remained non-turbid, including the controls (Fig. 5C). Parallel plating detected no pathogenic bacteria in these four specimens.

Observed growth after overnight incubation in GT-AST indicated levofloxacin susceptibility in all four culture-positive specimens from three patients, in agreement with the hospital clinical laboratory. Parallel plating and hospital testing also agreed on bacterial detection in all eight specimens from six patients (Supplementary Table S7).

At the 0.5 threshold, the model predicted no growth in some drug-exposed wells that grew overnight. All of these wells were from BALF001 (Fig. 5C). The MIC predicted from the concentration series for this specimen was ≤0.5 µg/mL in 63 evaluations (63%) and 1 µg/mL in 37 (37%).

The model predicted MICs for all four culture-positive BALF specimens in 96 of the 100 evaluations. Only BALF006 lacked a prediction in the remaining four. Mean MIC prediction coverage was 99.0%. Across the 100 evaluations, exact and essential agreement with hospital MICs also averaged 99.0% (Supplementary Table S7).

## 4. Discussion

GT-AST brings paired glucose-turbidity measurements, early growth prediction and direct specimen testing together to explore a route to earlier susceptibility information for antibiotic selection. Under the same antibiotic exposure conditions, glucose-associated and turbidity responses diverged in some exposed cultures: attenuated turbidity responses coexisted with glucose signals consistent with substrate utilization. Similar early separation was followed by either growth or no growth after overnight incubation (Figs. 2 and 3). Paired measurements therefore add information about responses during drug exposure to the later assessment of visible growth. Growth prediction and comparison with reference MICs connect these early responses to the growth-inhibition information used to guide treatment selection for patients.

This divergence is biologically plausible because substrate utilization and culture turbidity reflect different aspects of bacterial physiology. Continued glucose consumption after growth arrest and experimental separation of metabolic state from growth have been demonstrated in *other systems* (Chubukov and Sauer, 2014; Lopatkin et al., 2019). In GT-AST, declining residual-glucose signals alongside limited turbidity changes suggest that glucose-utilizing activity may continue during early drug exposure (Fig. 2). The distinction between activity and growth is also relevant to antibiotic persistence and VBNC states. Persister cells survive bactericidal exposure longer than most cells in the population and can regrow after drug removal, whereas VBNC cells remain viable but do not grow under the culture conditions examined (Balaban et al., 2019; Bamford et al., 2017). Paired measurements could help explore how substrate utilization relates to survival and recovery in these distinct states.

Growth prediction links the paired early responses to the growth inhibition assessed by phenotypic AST (Fig. 4). Conventional BMD relies on a growth reading after overnight incubation; GT-AST uses responses collected during approximately 2 h of antibiotic exposure to predict subsequent growth at each concentration. Across the 228 isolate-drug combinations, predicted MICs showed mean EA of 87.1% with independent reference MICs (Supplementary Table S4). These findings support using early paired responses to anticipate growth-inhibition endpoints. Further evaluation should examine the concentration-specific prediction errors that shift the MIC estimate and their distribution across isolate-drug combinations.

Direct BALF testing extends the paired-response approach to clinical specimens by starting measurements before bacterial isolation. Levofloxacin susceptibility classifications based on observed growth after overnight incubation agreed with the hospital clinical laboratory in all four culture-positive specimens. Parallel plating and hospital testing also agreed on bacterial detection in all eight specimens (Supplementary Table S7).

Across the four culture-positive specimens from three patients, model-predicted MICs showed mean EA of 99.0% with hospital MICs (Supplementary Table S7). In BALF001, the model missed growth in some drug-exposed wells, but both predicted MICs fell within the hospital-reported range. In BALF006, a missed growth call for the no-drug control prevented MIC estimation in four evaluations (Fig. 5). These observations highlight prediction accuracy and replicate concordance near the growth-inhibition transition as key requirements for translating early well-level predictions into reliable MIC estimates. Further validation should extend these paired MIC comparisons across a broader range of clinical specimens.

Direct gradient diffusion on selected bronchial aspirates and automated microscopy of mini-BAL samples have also explored susceptibility testing directly from respiratory specimens (Burillo et al., 2019; Douglas et al., 2015). In the retrospective input comparison, the model without species input produced BALF growth predictions using glucose and turbidity trajectories, drug and concentration inputs (Supplementary Table S8). This finding supports examining direct BALF workflows in which phenotypic assessment and species identification proceed in parallel. Combining these measurements with rapid species identification could bring together drug-response information and pathogen identity earlier, offering the potential to issue a complete microbiological report sooner.

Molecular and phenotypic tests provide complementary information for antibiotic selection. Molecular panels identify selected organisms and resistance genes, while conventional BMD and automated phenotypic systems assess growth inhibition during antibiotic exposure. Nanomotion sensing and measurements of extracellular metabolic changes extend the range of responses used for phenotypic AST (Sturm et al., 2024; Rydzak et al., 2022). GT-AST pairs residual-glucose signals with turbidity within the same concentration series, allowing glucose utilization to be considered alongside growth-related responses. Supplementary Table S5 compares these approaches by starting material, output and equipment requirements. Their clinical turnaround should be evaluated from specimen receipt to report completion, including preparation, identification and measurement.

GT-AST uses routine microbiology laboratory instruments, with estimated reagent costs of approximately US$0.50-0.60 per BALF specimen per experiment (Supplementary Table S6). These modest reagent requirements support the practical development of paired glucose-turbidity measurements for phenotypic AST.

Model development used 43 cultured isolates; direct BALF testing included eight specimens from six patients and focused on levofloxacin. A larger collection of paired responses and independently measured reference MICs should cover more isolate-drug combinations, clinical specimens and bacterial loads. Measurements near growth-inhibition transitions would help determine which concentration-specific errors alter MIC estimates and how consistently replicate wells resolve these transitions. Sharing these paired data across laboratories would support evaluation of reproducibility across operators, batches and laboratories. Automated liquid handling could reduce operator-dependent variation and handling time, with these effects assessed during workflow evaluation.

The separation of glucose and turbidity responses also raises a physiological question about the relationship between substrate utilization and bacterial survival. Larger collections of paired responses would help clarify the physiological significance of the patterns identified by exploratory screening. Combining paired trajectories with independent viability assays, time-kill measurements and regrowth after antibiotic removal would relate early glucose utilization to survival during exposure and subsequent recovery. Responses in independently characterized persister and VBNC states could then be compared to test whether they exhibit distinguishable paired patterns (Balaban et al., 2019; Bamford et al., 2017). Such comparisons would assess whether glucose measurements add information about survival and recovery to growth-based susceptibility assessment.

Our findings link paired glucose-turbidity responses to subsequent growth inhibition and extend MIC estimation to BALF specimens tested without prior bacterial isolation. Developing these early measurements alongside rapid pathogen identification could bring susceptibility information into treatment decisions sooner, helping patients receive effective and appropriately targeted antibiotics.

## Ethics statement

The Ethics Committee of Fuzhou University Affiliated Provincial Hospital issued ethics review reference K2023-08-018.

## CRediT authorship contribution statement

**Yuanzhuo Deng**: Writing - original draft, Visualization, Formal analysis, Data curation, Software, Investigation, Methodology. **Shaohan Xu**: Resources, Funding acquisition, Validation. **Huan Hu**: Supervision. **Haitao Yang**: Resources. **Wenqian Jiang**: Project administration, Funding acquisition, Methodology. **Tianzhu Zhu**: Software. **Qing Li**: Investigation. **Jun Lin**: Conceptualization, Funding acquisition, Methodology, Project administration, Resources, Supervision, Writing - review & editing.

## Funding

This work was supported by the Joint Funds for the Innovation of Science and Technology, Fujian Province (2024Y9060, 2024Y9570, and 2025Y9010), the Fujian Key Laboratory of Women and Children’s Critical Diseases Research (2024OP002), and the Fujian Provincial Natural Science Foundation (2026J0011679).

## Declaration of competing interests

A patent application related to this work is pending.

## Declaration of AI assistance

OpenAI Codex (GPT-6.0 Astra) was utilized to assist in translating the manuscript into English and refining the language for clarity. Additionally, it aided in revising the code for modeling and data analysis.

## References

Anderson, M.J., 2001. A new method for non-parametric multivariate analysis of variance. Austral Ecology 26, 32–46. 10.1111/j.1442-9993.2001.01070.pp.x

Balaban, N.Q., Helaine, S., Lewis, K., Ackermann, M., Aldridge, B., Andersson, D.I., Brynildsen, M.P., Bumann, D., Camilli, A., Collins, J.J., Dehio, C., Fortune, S., Ghigo, J.M., Hardt, W.D., Harms, A., Heinemann, M., Hung, D.T., Jenal, U., Levin, B.R., Michiels, J., Storz, G., Tan, M.W., Tenson, T., Van Melderen, L., Zinkernagel, A., 2019. Definitions and guidelines for research on antibiotic persistence. Nature Reviews Microbiology 17, 441–448. 10.1038/s41579-019-0196-3

Baltekin, Ö., Boucharin, A., Tano, E., Andersson, D.I., Elf, J., 2017. Antibiotic susceptibility testing in less than 30 min using direct single-cell imaging. Proceedings of the National Academy of Sciences 114, 9170–9175. 10.1073/pnas.1708558114

Bamford, R.A., Smith, A., Metz, J., Glover, G., Titball, R.W., Pagliara, S., 2017. Investigating the physiology of viable but non-culturable bacteria by microfluidics and time-lapse microscopy. BMC Biology 15, 121. 10.1186/s12915-017-0465-4

bioMérieux, n.d.-a. BIOFIRE Blood Culture Identification 2 (BCID2) Panel. https://www.biomerieux.com/us/en/our-offer/clinical-products/biofire-blood-culture-identification-2-panel.html (accessed 5 October 2026).

bioMérieux, n.d.-b. VITEK REVEAL: Fast antimicrobial susceptibility testing. https://www.biomerieux.com/corp/en/our-offer/clinical-products/vitek-reveal.html (accessed 5 October 2026).

Burillo, A., de Egea, V., Onori, R., Martín-Rabadán, P., Cercenado, E., Jiménez-Navarro, L., Muñoz, P., Bouza, E., 2019. Gradient diffusion antibiogram used directly on bronchial aspirates for a rapid diagnosis of ventilator-associated pneumonia. Antimicrobial Resistance & Infection Control 8, 176. 10.1186/s13756-019-0640-1

Chubukov, V., Sauer, U., 2014. Environmental Dependence of Stationary-Phase Metabolism in *Bacillus subtilis* and *Escherichia coli*. Applied and Environmental Microbiology 80, 2901–2909. 10.1128/aem.00061-14

Clinical and Laboratory Standards Institute (CLSI), 2023. Breakpoint Implementation Toolkit: Accuracy Instructions for Prefilled Excel (Part E), version 1.0. https://clsi.org/media/i3ylyrde/part_e_accuracy_instructions_for_prefilled_excel_workbook.pdf (accessed 25 October 2026).

Clinical and Laboratory Standards Institute (CLSI), 2026. Performance Standards for Antimicrobial Susceptibility Testing. 36th ed. CLSI supplement M100. Clinical and Laboratory Standards Institute. https://clsi.org/shop/standards/m100/ (accessed 25 October 2026).

Douglas, I.S., Price, C.S., Overdier, K.H., Wolken, R.F., Metzger, S.W., Hance, K.R., Howson, D.C., 2015. Rapid Automated Microscopy for Microbiological Surveillance of Ventilator-associated Pneumonia. American Journal of Respiratory and Critical Care Medicine 191, 566–573. 10.1164/rccm.201408-1468oc

EUCAST, 2024. Reading guide for broth microdilution, version 5.0. https://www.eucast.org/fileadmin/eucast/pdf/MIC/Reading_guide_BMD_v_5.0_2024.pdf (accessed 30 September 2026).

EUCAST, 2026. Breakpoint tables for interpretation of MICs and zone diameters, version 16.1. https://www.eucast.org/fileadmin/eucast/pdf/breakpoints/v_16.1_Breakpoint_Tables.pdf (accessed 23 October 2026).

Gower, J.C., 1966. Some distance properties of latent root and vector methods used in multivariate analysis. Biometrika 53, 325–338. 10.1093/biomet/53.3-4.325

Hultman, E., 1959. Rapid Specific Method for Determination of Aldosaccharides in Body Fluids. Nature 183, 108–109. 10.1038/183108a0

International Organization for Standardization (ISO), 2021. ISO 20776-2:2021. Clinical laboratory testing and in vitro diagnostic test systems — Susceptibility testing of infectious agents and evaluation of performance of antimicrobial susceptibility test devices — Part 2: Evaluation of performance of antimicrobial susceptibility test devices against reference broth micro-dilution. 2nd ed. https://www.iso.org/standard/79377.html (accessed 25 October 2026).

Jafari, M.J., Golabi, M., Ederth, T., 2024. Antimicrobial susceptibility testing using infrared attenuated total reflection (IR-ATR) spectroscopy to monitor metabolic activity. Spectrochimica Acta Part A: Molecular and Biomolecular Spectroscopy 304, 123384. 10.1016/j.saa.2023.123384

Loiodice, A., Bailly, S., Ruckly, S., Buetti, N., Barbier, F., Staiquly, Q., Tabah, A., Timsit, J.-F., on behalf of the EUROBACT-2 Study Group, ESICM, ESGCIP and the OUTCOMEREA Network, 2024. Effect of adequacy of empirical antibiotic therapy for hospital-acquired bloodstream infections on intensive care unit patient prognosis: a causal inference approach using data from the Eurobact2 study. Clinical Microbiology and Infection 30, 1559–1568. 10.1016/j.cmi.2024.09.011

Lopatkin, A.J., Stokes, J.M., Zheng, E.J., Yang, J.H., Takahashi, M.K., You, L., Collins, J.J., 2019. Bacterial metabolic state more accurately predicts antibiotic lethality than growth rate. Nature Microbiology 4, 2109–2117. 10.1038/s41564-019-0536-0

Loshchilov, I., Hutter, F., 2019. Decoupled Weight Decay Regularization. International Conference on Learning Representations. https://arxiv.org/abs/1711.05101

Paszke, A., Gross, S., Massa, F., Lerer, A., Bradbury, J., Chanan, G., Killeen, T., Lin, Z., Gimelshein, N., Antiga, L., Desmaison, A., Köpf, A., Yang, E., DeVito, Z., Raison, M., Tejani, A., Chilamkurthy, S., Steiner, B., Fang, L., Bai, J., Chintala, S., 2019. PyTorch: An Imperative Style, High-Performance Deep Learning Library. Advances in Neural Information Processing Systems 32. https://papers.nips.cc/paper_files/paper/2019/hash/bdbca288fee7f92f2bfa9f7012727740-Abstract.html

Pedregosa, F., Varoquaux, G., Gramfort, A., Michel, V., Thirion, B., Grisel, O., Blondel, M., Prettenhofer, P., Weiss, R., Dubourg, V., Vanderplas, J., Passos, A., Cournapeau, D., Brucher, M., Perrot, M., Duchesnay, É., 2011. Scikit-learn: Machine Learning in Python. Journal of Machine Learning Research 12, 2825–2830. https://jmlr.org/papers/v12/pedregosa11a.html

Riester, O., Kaiser, L., Laufer, S., Deigner, H.P., 2024. Rapid Phenotypic Antibiotics Susceptibility Analysis by a 3D Printed Prototype. Advanced Science 11, 2308806. 10.1002/advs.202308806

Rydzak, T., Groves, R.A., Zhang, R., Aburashed, R., Pushpker, R., Mapar, M., Lewis, I.A., 2022. Metabolic preference assay for rapid diagnosis of bloodstream infections. Nature Communications 13, 2332. 10.1038/s41467-022-30048-6

Sturm, A., Jóźwiak, G., Verge, M.P., Munch, L., Cathomen, G., Vocat, A., Luraschi-Eggemann, A., Orlando, C., Fromm, K., Delarze, E., Świątkowski, M., Wielgoszewski, G., Totu, R.M., García-Castillo, M., Delfino, A., Tagini, F., Kasas, S., Lass-Flörl, C., Gstir, R., Cantón, R., Greub, G., Cichocka, D., 2024. Accurate and rapid antibiotic susceptibility testing using a machine learning-assisted nanomotion technology platform. Nature Communications 15, 2037. 10.1038/s41467-024-46213-y

U.S. Food and Drug Administration, 2019. VITEK 2 AST - Gram Negative Imipenem: K183415. https://www.accessdata.fda.gov/cdrh_docs/pdf18/K183415.pdf (accessed 21 October 2026).

U.S. Food and Drug Administration, 2020. BioFire Blood Culture Identification 2 (BCID2) Panel: 510(k) substantial equivalence determination decision summary, K193519. https://www.accessdata.fda.gov/cdrh_docs/reviews/K193519.pdf (accessed 5 October 2026).

Zhang, M., Hong, W., Abutaleb, N.S., Li, J., Dong, P.T., Zong, C., Wang, P., Seleem, M.N., Cheng, J.X., 2020. Rapid Determination of Antimicrobial Susceptibility by Stimulated Raman Scattering Imaging of D2O Metabolic Incorporation in a Single Bacterium. Advanced Science 7, 2001452. 10.1002/advs.202001452

