## Supplementary Information for "Linking Early Bacterial Responses to Growth Inhibition via Glucose-Turbidity Dynamics for Rapid Antimicrobial Susceptibility Testing"

Contents

**Supplementary Figures S1-S9	27**

Supplementary Figure S1	27

Supplementary Figure S2	28

Supplementary Figure S3	29

Supplementary Figure S4	30

Supplementary Figure S5	31

Supplementary Figure S6	32

Supplementary Figure S7	34

Supplementary Figure S8	36

Supplementary Figure S9	37

**Supplementary Tables S1-S9	38**

Supplementary Table S1	38

Supplementary Table S2	39

Supplementary Table S3	40

Supplementary Table S4	42

Supplementary Table S5	43

Supplementary Table S6	45

Supplementary Table S7	46

Supplementary Table S8	47

Supplementary Table S9	51

**Supplementary Text	52**

Methodology	52

Results	56

**Supplementary References	57**

Supplementary Figures S1-S9

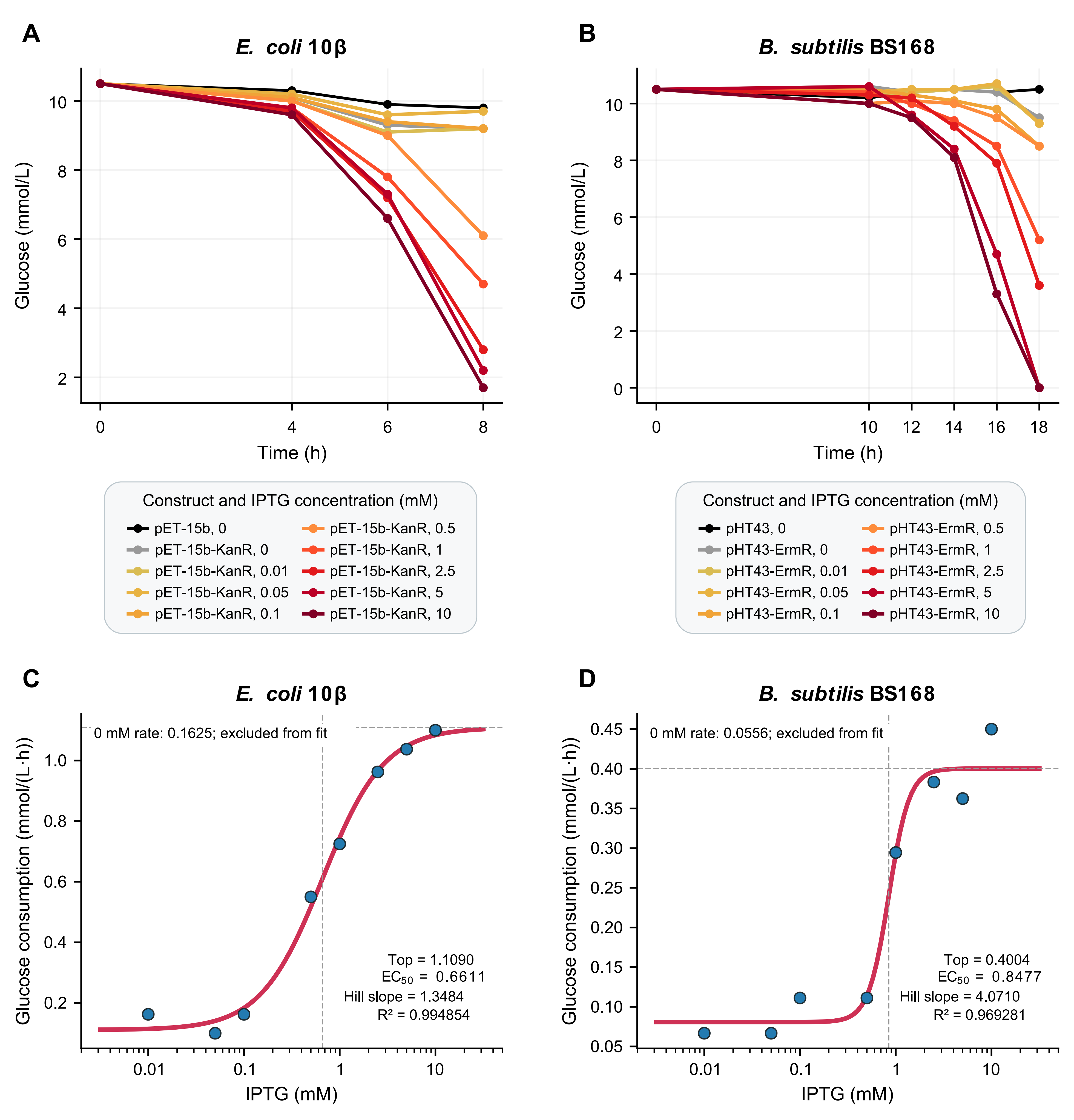

Supplementary Figure S1. Glucose consumption by engineered bacteria across IPTG concentrations

A, Glucose concentration time courses for *Escherichia coli* 10β carrying pET-15b or pET-15b-KanR in 10 µg/mL kanamycin. B, Corresponding time courses for *Bacillus subtilis* BS168 carrying pHT43 or pHT43-ErmR in 50 µg/mL erythromycin. Recombinant strains were tested at IPTG concentrations of 0-10 mM, and empty-vector controls received 0 mM IPTG. *E. coli* was precultured overnight in liquid medium containing the IPTG concentration used in the subsequent exposure condition. Points show condition means from a single experiment for each model, lines connect successive time points, and colours identify the conditions in the legends.

C and D, Average glucose consumption rates of the recombinant *E. coli* and *B. subtilis* strains in A and B, respectively. Blue points show rates calculated from the mean time courses between the initial and last strictly positive glucose readings; red curves show four-parameter Hill fits to positive IPTG concentrations. Positive concentrations are displayed on a logarithmic axis; the 0 mM rates are listed separately within the panels and are excluded from fitting. Horizontal and vertical dashed lines indicate the fitted Top and EC_50_, respectively. IPTG, isopropyl β-D-1-thiogalactopyranoside. Bottom and Top are the fitted lower and upper plateaus; EC_50_ is the IPTG concentration at the midpoint between them, in mM.

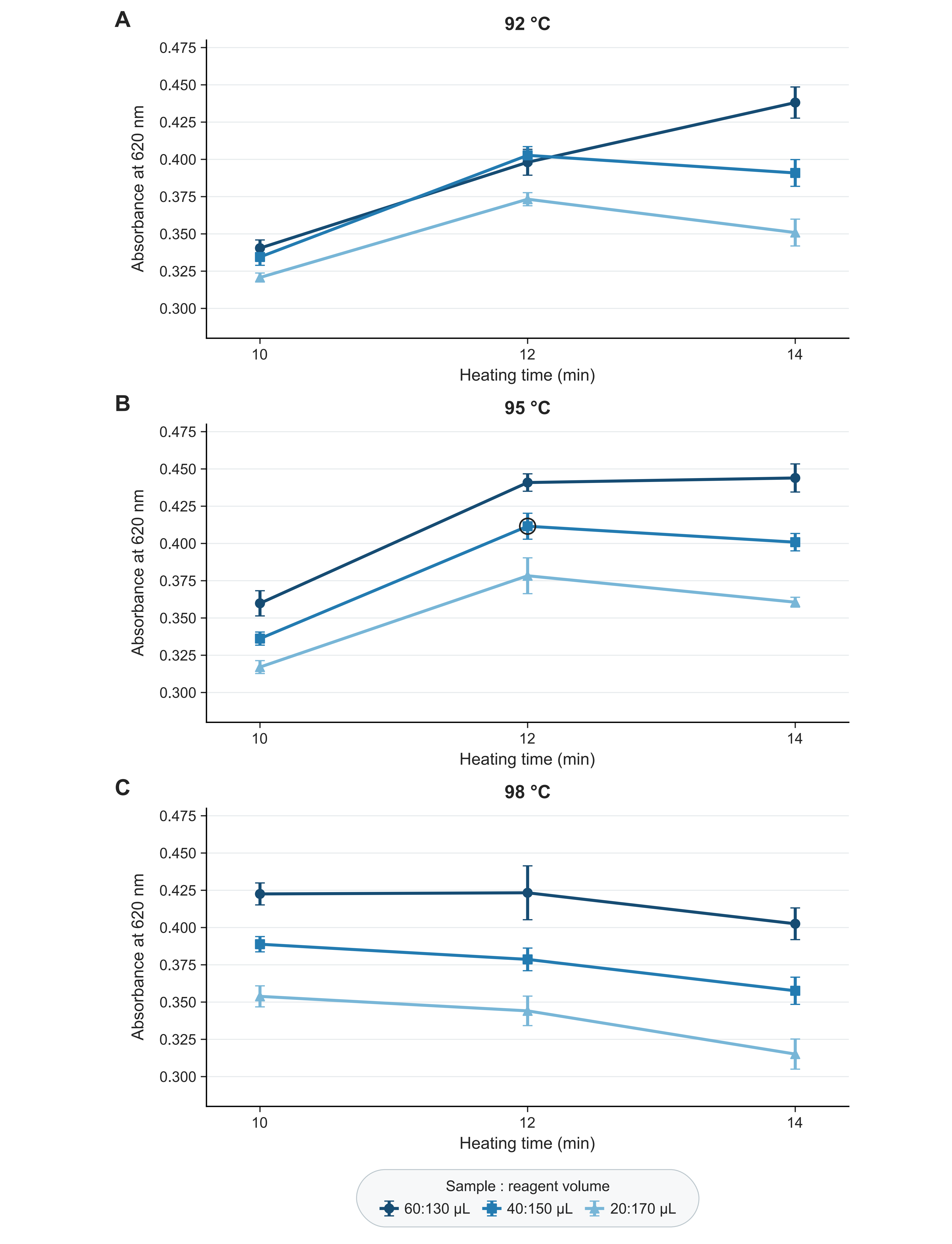

Supplementary Figure S2. Glucose color development across heating and volume conditions

A-C, Absorbance at 620 nm for 400 mg/dL glucose in MH medium heated at 92, 95 and 98 °C, respectively. Dark-blue circles, medium-blue squares and light-blue triangles identify sample:o-toluidine reagent volumes of 60:130, 40:150 and 20:170 µL, respectively. Points and error bars show the mean and sample standard deviation (SD) of eight technical replicate wells per condition; lines connect the measurements at 10, 12 and 14 min. The black ring marks the selected conditions of 95 °C, 12 min and 40:150 µL. Summary statistics are provided in Supplementary Table S1. MH, Mueller-Hinton.

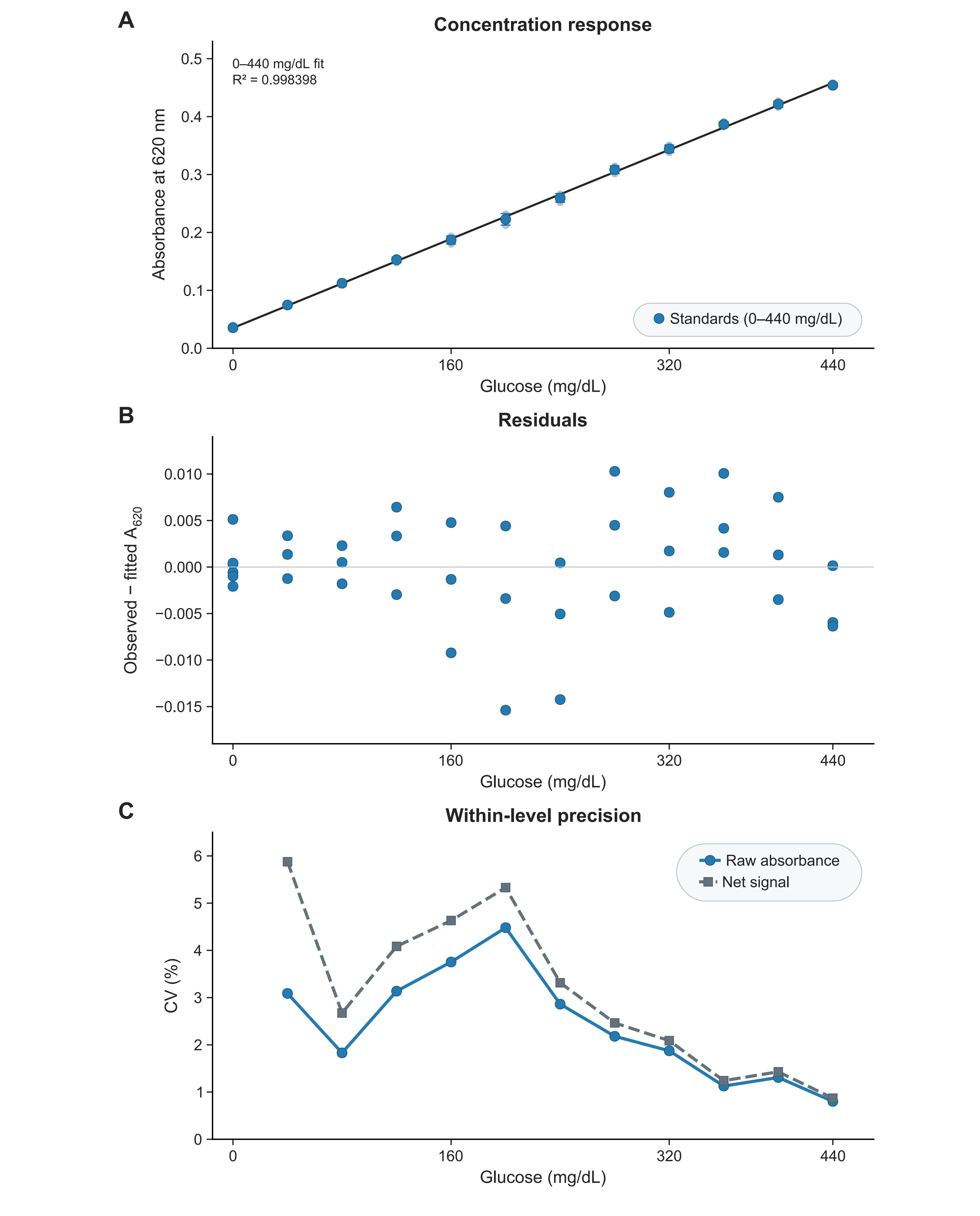

Supplementary Figure S3. Glucose calibration and precision in MH medium

Glucose standards in MH medium were heated at 95 °C for 12 min with 40 µL sample and 150 µL o-toluidine reagent. A, Absorbance at 620 nm over 0-440 mg/dL. Blue points and error bars show the mean and sample standard deviation (SD), and pale points show individual wells. The black line is an ordinary-least-squares fit to all 39 wells, with equal weight per well and a freely estimated intercept. B, Individual-well residuals (observed minus fitted absorbance); the horizontal line marks zero. C, Coefficients of variation (CV) for nonzero standards (40-440 mg/dL). Blue circles and solid lines show raw-absorbance CV; gray squares and dashed lines show net-signal CV after subtraction of the mean of the six zero-concentration MH wells. The zero-concentration standard has six technical replicate wells, and each nonzero concentration has three. Concentrations refer to the MH solution before mixing with reagent. Summary statistics and fit parameters are provided in Supplementary Table S2. MH, Mueller-Hinton.

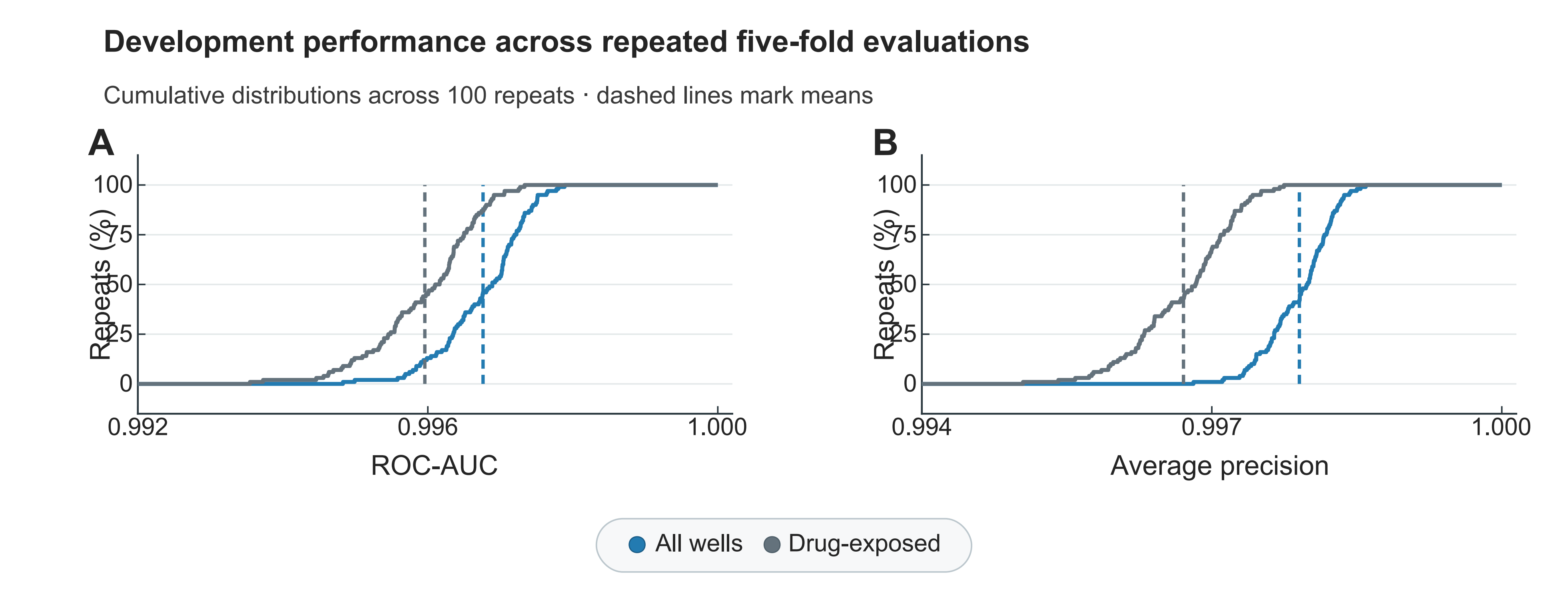

Supplementary Figure S4. Performance of the full model across 100 repeats of isolate-grouped five-fold cross-validation

A, ROC-AUC across 100 evaluations of the full model. B, Corresponding average precision. Blue and gray curves show cumulative distributions for all wells and drug-exposed wells, respectively. Dashed lines mark the means. Each evaluation pools the predictions for held-out isolates across folds. Growth labels were assigned by manual assessment of visible turbidity after overnight incubation. Mean ROC and precision-recall curves are shown in Fig. 4A. T1, first reading; ROC-AUC, area under the receiver operating characteristic curve.

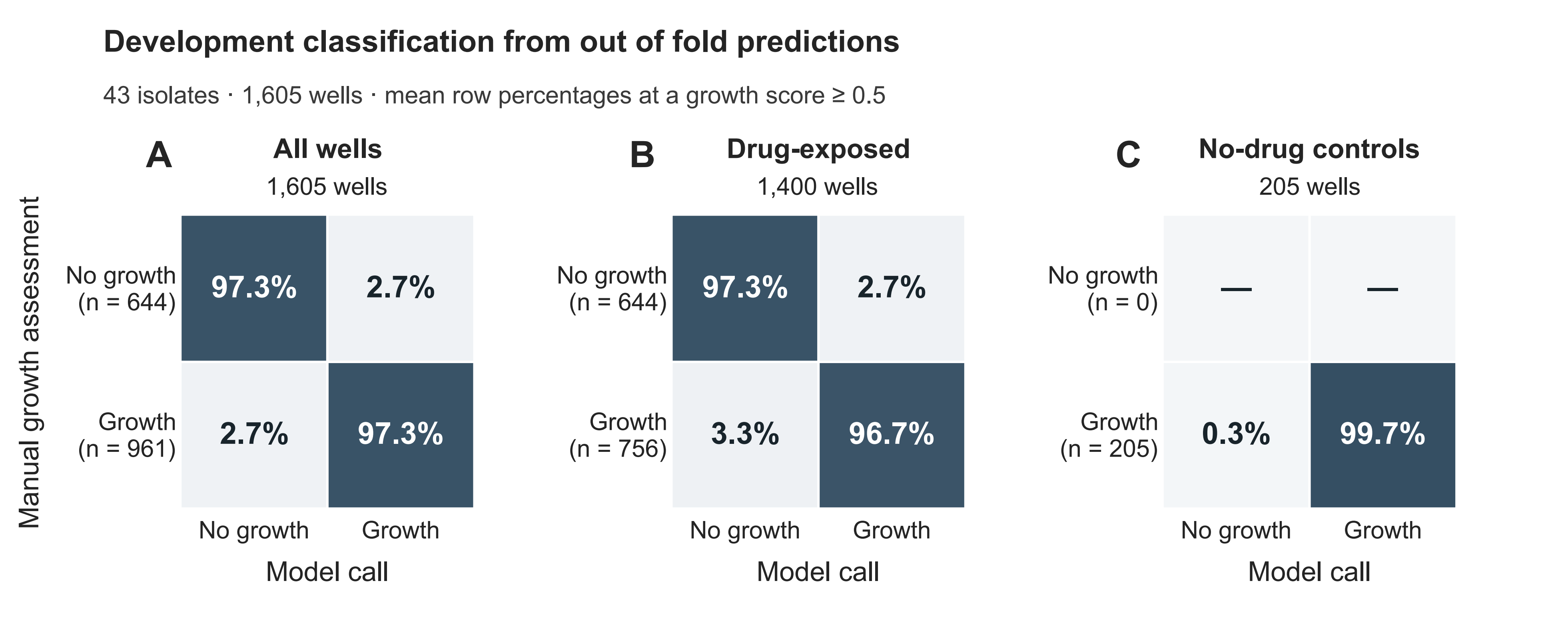

Supplementary Figure S5. Development confusion matrices for the full model

The matrices summarize out of fold predictions from the 100 isolate-grouped five-fold evaluations in Fig. 4A, covering 1,605 wells from 43 cultured isolates. Scores ≥0.5 predict growth. Growth labels were assigned by manual assessment of visible turbidity after overnight incubation. Each well was scored by models that excluded its isolate from training and early stopping.

A, All wells (1,605). B, Drug-exposed wells (1,400). C, No-drug controls (205). Rows show manually assessed growth labels and columns show model predictions. Across all wells, the observed outcomes were growth (n = 961, 59.9%) and no growth (n = 644, 40.1%). Among drug-exposed wells, the corresponding counts were 756 (54%) and 644 (46%). All 205 no-drug controls were labeled as growth, leaving no observations for estimating specificity.

Cell values and blue-gray shading show the mean percentages within each row. TP, true positive; FN, false negative; FP, false positive; TN, true negative. Growth assessed manually after overnight incubation is the positive reference class. T1, first reading.

Supplementary Figure S6

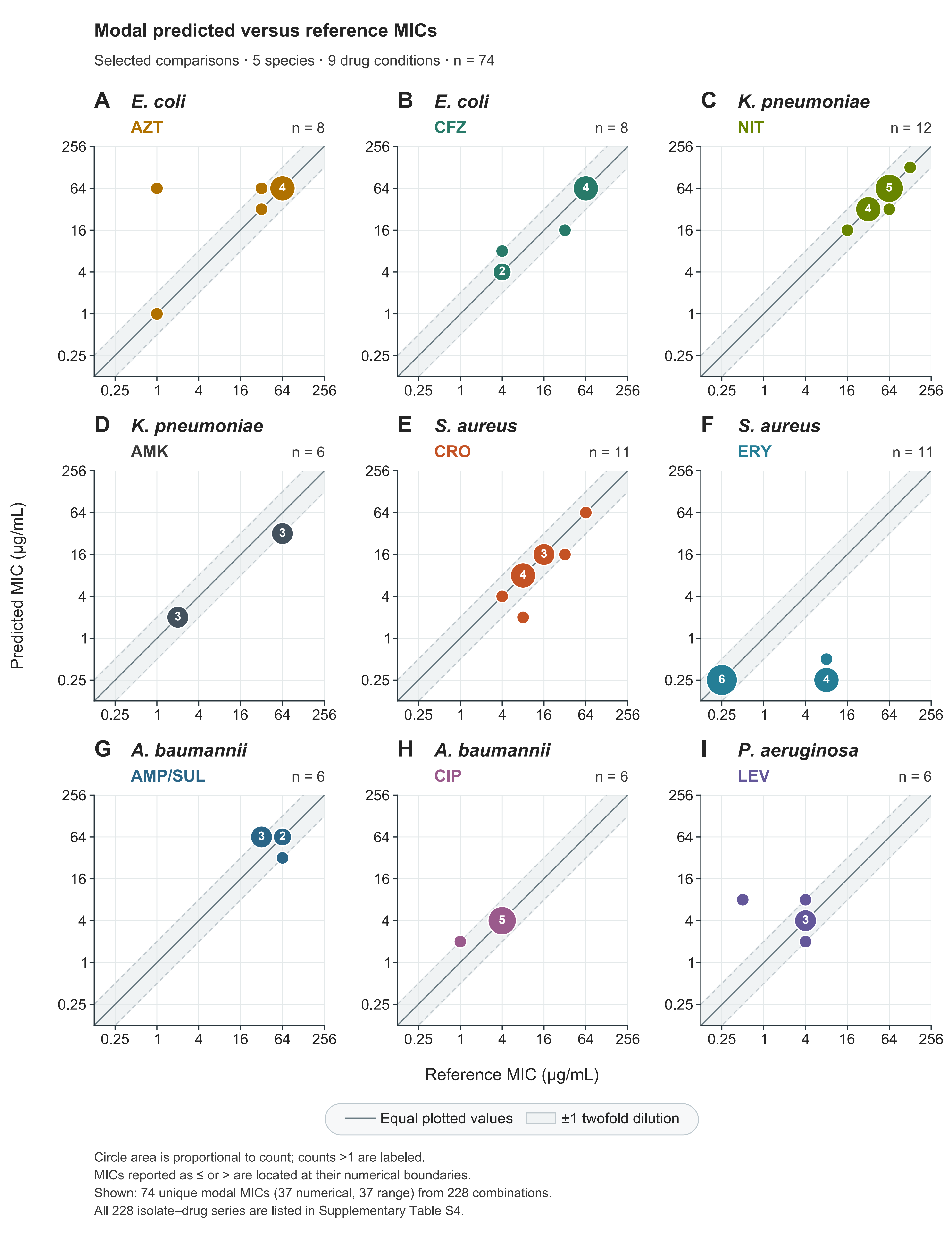

Supplementary Figure S6. Agreement between predicted and reference MICs

Each panel compares the most frequent predicted MIC across 100 repeats with the reference MIC for a selected species-drug combination, with 74 comparisons in total. A-B, *E. coli* with AZT (n = 8) and CFZ (n = 8). C-D, *K. pneumoniae* with NIT (n = 12) and AMK (n = 6). E-F, *S. aureus* with CRO (n = 11) and ERY (n = 11). G-H, *A. baumannii* with AMP/SUL (n = 6) and CIP (n = 6). I, *P. aeruginosa* with LEV (n = 6). MICs reported with ≤ or > are plotted at the stated numerical bounds. Circle colors distinguish drugs, and circle area is proportional to the number of comparisons at the same coordinates; counts greater than one are shown inside circles. Both axes use log₂ scales. The solid line marks equality; dashed lines and shading mark ±1 twofold dilution. For AMP/SUL, both axes use the ampicillin concentration. Supplementary Table S4 lists all 228 isolate-drug combinations. Across repeats, mean exact agreement was 64.8% and mean essential agreement (EA) was 87.1% under the comparison rules in Methods 2.14. MIC, minimum inhibitory concentration; AMK, amikacin; AMP/SUL, ampicillin-sulbactam; AZT, aztreonam; CFZ, cefazolin; CIP, ciprofloxacin; CRO, ceftriaxone; ERY, erythromycin; LEV, levofloxacin; NIT, nitrofurantoin.

Supplementary Figure S7

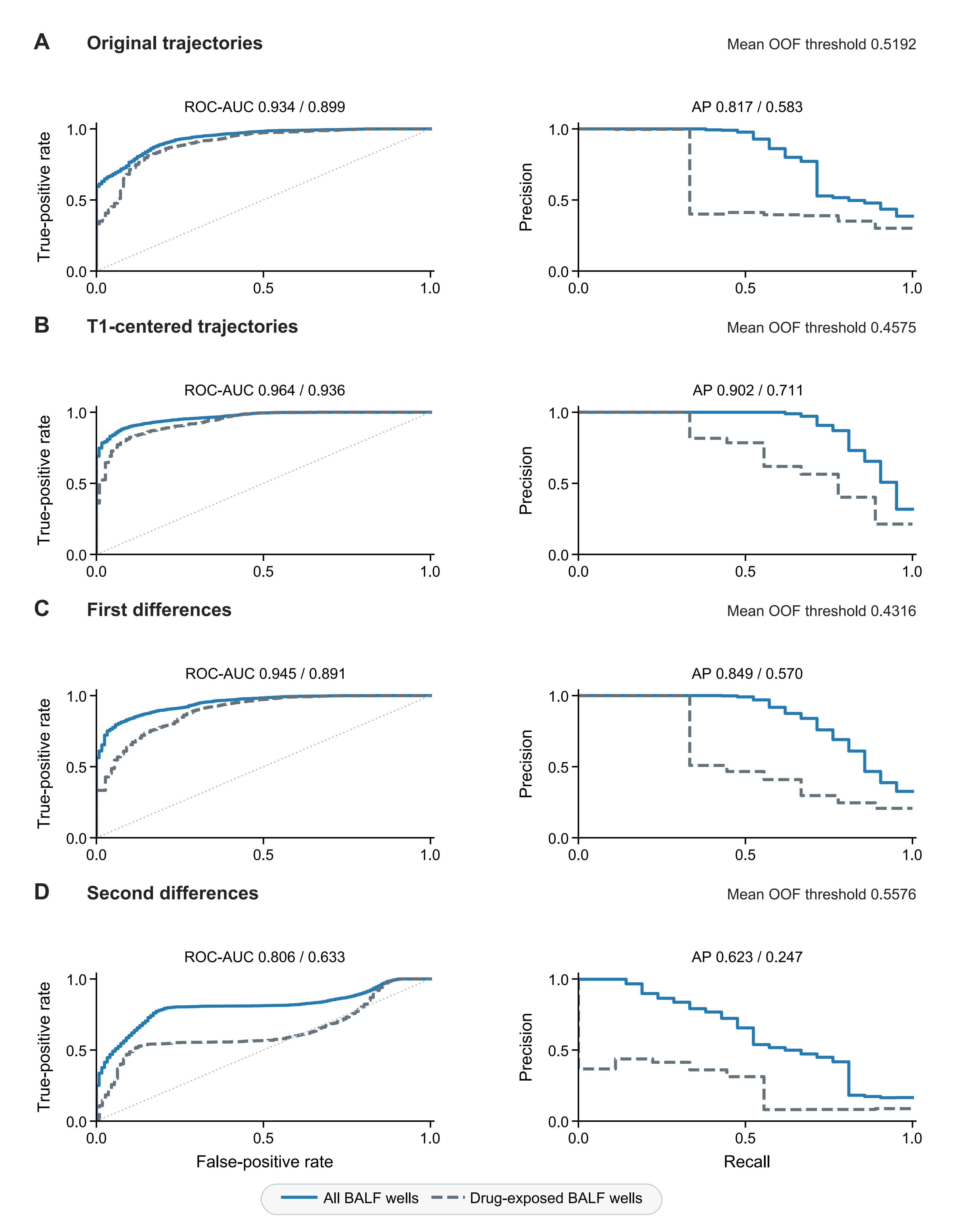

Supplementary Figure S7. Growth prediction from original, T1-centered and derivative trajectories

A-D show models using original trajectories, T1-centered trajectories, first differences and second differences, respectively. Each model combines glucose and turbidity inputs with species, drug and concentration features. Eight readings span 30-135 min at 15-min intervals, giving eight values per channel for original and centered trajectories, seven for first differences and six for second differences. Differences are divided by 15 min at each order. Subtracting T1 leaves both difference orders unchanged, so each is shown once.

Within each row, left and right plots show BALF ROC and precision-recall curves. Curves and metrics are means across 100 repeats. Blue solid lines represent all 144 wells (21 labeled as growth), and gray dashed lines represent 120 drug-exposed wells (9 labeled as growth); metric pairs above the plots follow this order. Growth labels were assigned by manual assessment of visible turbidity after overnight incubation. ROC dotted lines indicate random ranking. Pooled development out of fold metrics and BALF classification counts are provided in Supplementary Table S3. Models used the same isolate-grouped splits and training protocol; scaling used only optimization data.

Thresholds were selected from pooled development out of fold (OOF) scores in each repeat. Their means are displayed to four decimal places: 0.5192 in A, 0.4575 in B, 0.4316 in C and 0.5576 in D. BALF calls use the full-precision threshold from the corresponding repeat. Supplementary Table S3 compares performance at these thresholds and at the uniform threshold of 0.5000. The accompanying Table_S3_model_metrics.tsv gives the minimum and maximum TP, FN, FP and TN counts across repeats. ROC-AUC and AP use continuous scores and are identical under the two threshold rules.

BALF performance was evaluated retrospectively in eight specimens from six patients, with technical replicate wells nested within specimens. Model growth scores are uncalibrated. T1, first reading; BALF, bronchoalveolar lavage fluid. ROC, receiver operating characteristic; AUC, area under the ROC curve; AP, average precision. TP, true positive; FN, false negative; FP, false positive; TN, true negative. Growth assessed manually after overnight incubation is the positive reference class.

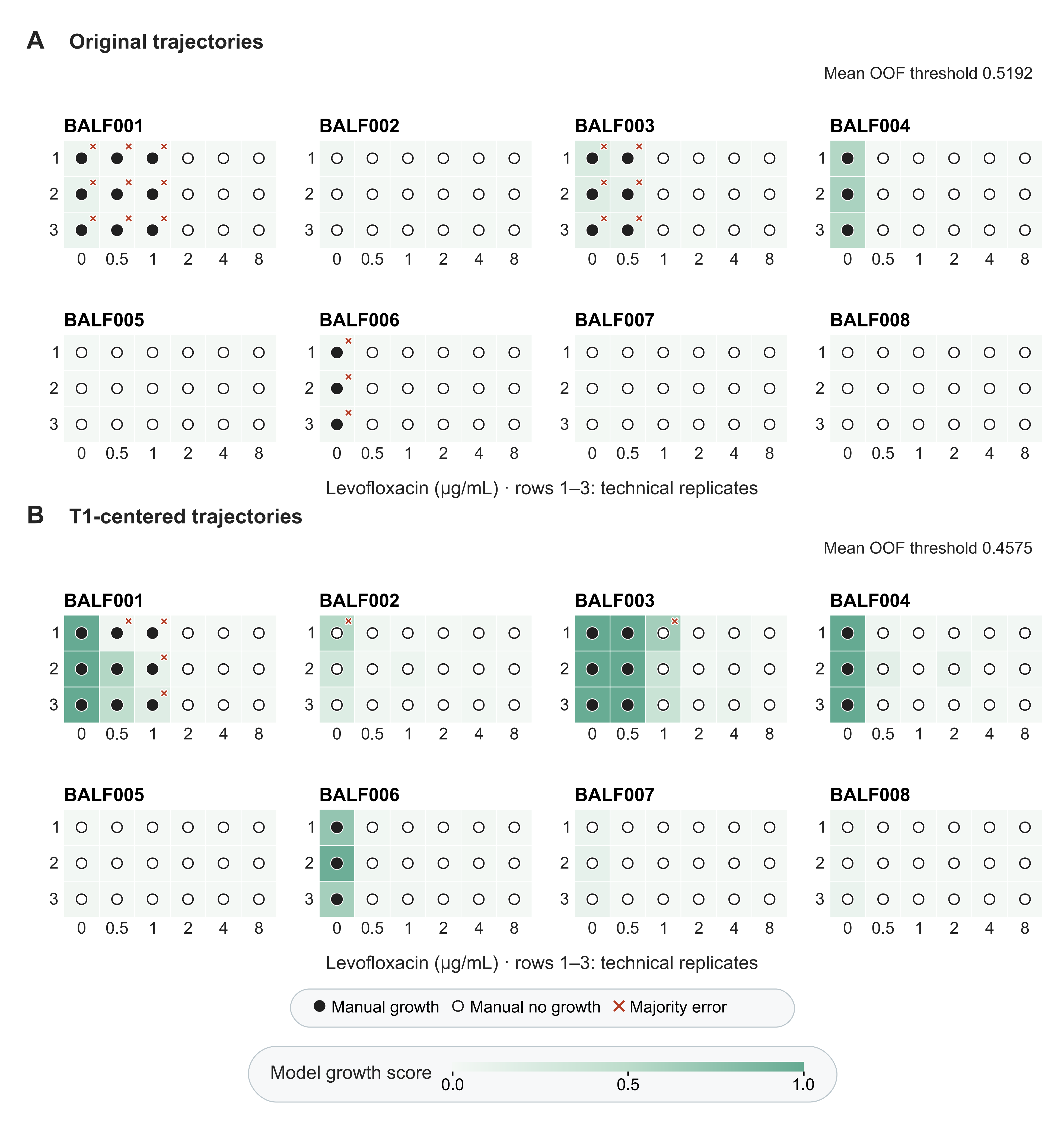

Supplementary Figure S8. Per-well BALF growth predictions from original and T1-centered trajectories

A, Predictions from the original paired glucose and turbidity trajectories. B, Predictions after subtracting each well's first reading (T1) from each channel, using its threshold selected from development out of fold (OOF) scores. The input trajectories comprise eight readings at 30-135 min after inoculation. Each panel shows 144 wells from eight bronchoalveolar lavage fluid (BALF) specimens. Columns show five levofloxacin (LEV) concentrations (0.5, 1, 2, 4 and 8 µg/mL) and the no-drug control (0); rows show three technical replicate wells per concentration.

Green shading indicates the mean uncalibrated model growth score (0-1) across 100 repeats. Solid black circles and white circles with black outlines indicate growth and no growth, respectively, assessed manually after overnight incubation. Growth is predicted when the score meets or exceeds the development OOF-selected threshold for that repeat. Panels display the mean threshold. Orange-red crosses mark wells that disagreed with the manual assessment in more than half of the 100 repeats. Of the 144 wells, crosses mark discrepant wells in A (n = 18, 12.5%) and B (n = 6, 4.2%).

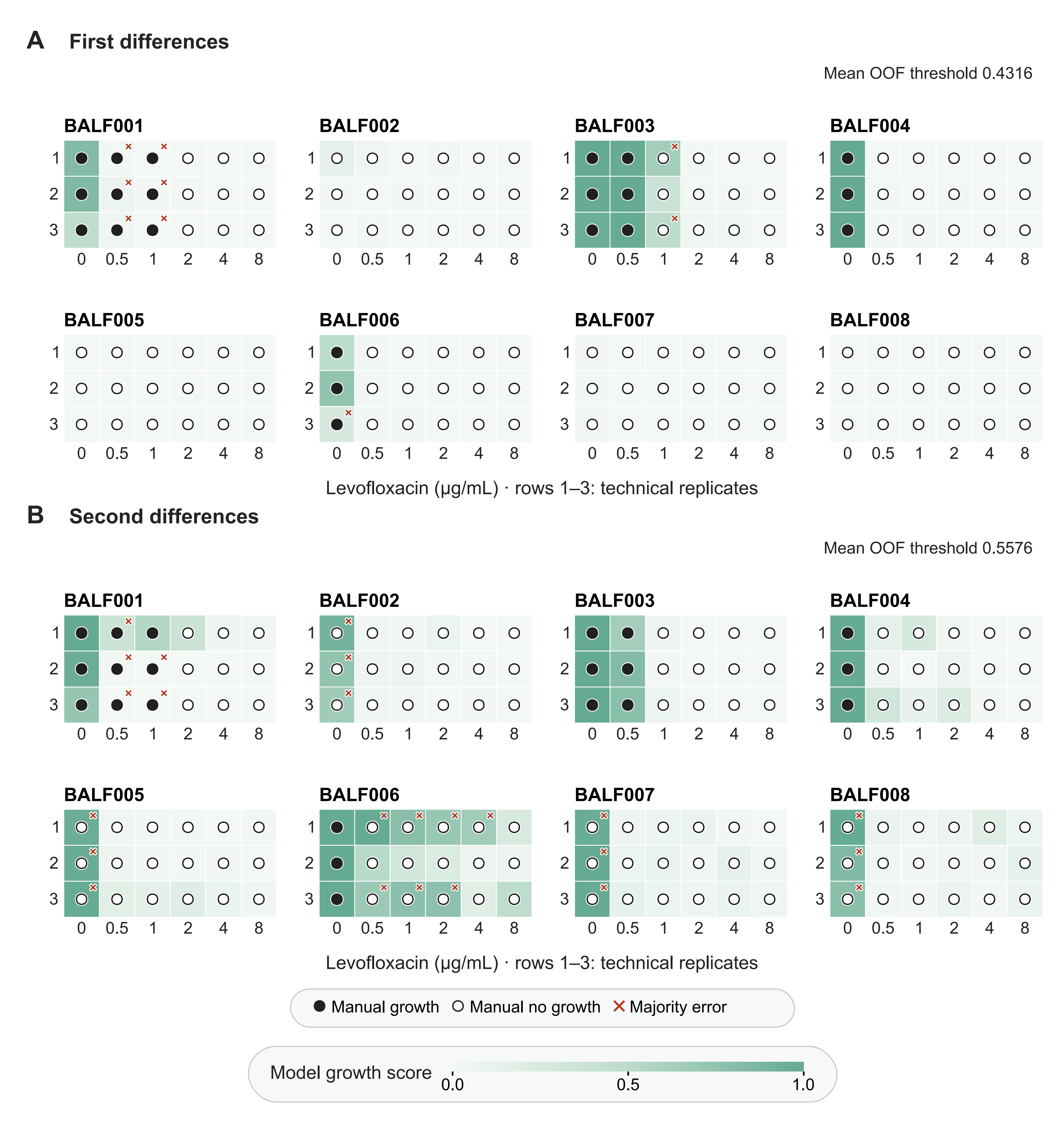

Supplementary Figure S9. Per-well BALF growth predictions from first and second differences

A, Predictions from first differences of the paired glucose and turbidity trajectories. B, Predictions from second differences. Both panels show 144 wells from eight bronchoalveolar lavage fluid (BALF) specimens and retain the concentration layout and well ordering of Supplementary Fig. S8.

Green shading indicates the mean uncalibrated model growth score (0-1) across 100 repeats. Solid black circles and white circles with black outlines indicate growth and no growth, respectively, assessed manually after overnight incubation. Orange-red crosses mark wells that disagreed with manual assessment in more than half of the 100 repeats at the repeat-specific development out of fold (OOF) thresholds (mean thresholds of 0.4316 in A and 0.5576 in B). Of the 144 wells, crosses mark discrepant wells in A (n = 9, 6.3%) and B (n = 24, 16.7%). Subtracting the first reading (T1) leaves both difference orders unchanged; the equivalent centered-input results are represented by these same panels.

Supplementary tables S1-S9

Supplementary Table S1. Glucose color development at 400 mg/dL

Each row summarizes one combination of heating temperature, heating time and sample:o-toluidine reagent volumes at 400 mg/dL glucose. The number of technical replicate wells is denoted by n (eight per condition). A_620_ denotes absorbance at 620 nm, reported as mean ± sample standard deviation (SD); CV is the coefficient of variation of raw absorbance. The selected conditions were 95 °C, 12 min and 40:150 µL.

| Temperature (°C) | Time (min) | Sample:reagent (µL) | n | A_620_ mean ± SD | CV (%) |
| --- | --- | --- | --- | --- | --- |
| 92 | 10 | 20:170 | 8 | 0.32075 ± 0.00300 | 0.94 |
| 92 | 10 | 40:150 | 8 | 0.33450 ± 0.00557 | 1.67 |
| 92 | 10 | 60:130 | 8 | 0.34045 ± 0.00548 | 1.61 |
| 92 | 12 | 20:170 | 8 | 0.37328 ± 0.00439 | 1.18 |
| 92 | 12 | 40:150 | 8 | 0.40272 ± 0.00584 | 1.45 |
| 92 | 12 | 60:130 | 8 | 0.39806 ± 0.00868 | 2.18 |
| 92 | 14 | 20:170 | 8 | 0.35090 ± 0.00900 | 2.56 |
| 92 | 14 | 40:150 | 8 | 0.39089 ± 0.00898 | 2.30 |
| 92 | 14 | 60:130 | 8 | 0.43811 ± 0.01046 | 2.39 |
| 95 | 10 | 20:170 | 8 | 0.31706 ± 0.00433 | 1.37 |
| 95 | 10 | 40:150 | 8 | 0.33618 ± 0.00441 | 1.31 |
| 95 | 10 | 60:130 | 8 | 0.35984 ± 0.00845 | 2.35 |
| 95 | 12 | 20:170 | 8 | 0.37834 ± 0.01202 | 3.18 |
| 95 | 12 | 40:150 | 8 | 0.41156 ± 0.00871 | 2.12 |
| 95 | 12 | 60:130 | 8 | 0.44088 ± 0.00584 | 1.32 |
| 95 | 14 | 20:170 | 8 | 0.36060 ± 0.00331 | 0.92 |
| 95 | 14 | 40:150 | 8 | 0.40086 ± 0.00583 | 1.45 |
| 95 | 14 | 60:130 | 8 | 0.44394 ± 0.00943 | 2.12 |
| 98 | 10 | 20:170 | 8 | 0.35384 ± 0.00705 | 1.99 |
| 98 | 10 | 40:150 | 8 | 0.38882 ± 0.00513 | 1.32 |
| 98 | 10 | 60:130 | 8 | 0.42256 ± 0.00737 | 1.74 |
| 98 | 12 | 20:170 | 8 | 0.34411 ± 0.00989 | 2.87 |
| 98 | 12 | 40:150 | 8 | 0.37864 ± 0.00760 | 2.01 |
| 98 | 12 | 60:130 | 8 | 0.42333 ± 0.01807 | 4.27 |
| 98 | 14 | 20:170 | 8 | 0.31511 ± 0.01010 | 3.21 |
| 98 | 14 | 40:150 | 8 | 0.35759 ± 0.00913 | 2.55 |
| 98 | 14 | 60:130 | 8 | 0.40254 ± 0.01063 | 2.64 |

Supplementary Table S2. Glucose calibration statistics

A, Summary statistics for glucose standards measured at 95 °C for 12 min with 40 µL sample and 150 µL o-toluidine reagent. C is the concentration in MH medium before mixing with reagent, and n is the number of technical replicate wells. A_620_ denotes absorbance at 620 nm, reported as mean ± sample standard deviation (SD). Raw CV is calculated from raw absorbance; Net CV uses the mean absorbance after subtraction of the mean of the six zero-concentration MH wells. Back-calculated C is derived from the mean absorbance at each concentration; deviation is its relative difference from nominal C. Dashes denote undefined net-signal CV and relative deviation at 0 mg/dL. B, Ordinary-least-squares calibration over 0-440 mg/dL, using all 39 individual wells with equal weight and a freely estimated intercept. R² and residual SD refer to this fit; the lack-of-fit p value uses pure error estimated from replicate wells at each concentration. CV, coefficient of variation; MH, Mueller-Hinton.

A Concentration-level summaries

| C (mg/dL) | n | A_620_ mean ± SD | Raw CV (%) | Net CV (%) | Back-calculated C (mg/dL) | Deviation (%) |
| --- | --- | --- | --- | --- | --- | --- |
| 0 | 6 | 0.03545 ± 0.00250 | 7.06 | — | 0.39 | — |
| 40 | 3 | 0.07470 ± 0.00231 | 3.09 | 5.88 | 41.21 | +3.01 |
| 80 | 3 | 0.11233 ± 0.00206 | 1.83 | 2.67 | 80.34 | +0.43 |
| 120 | 3 | 0.15273 ± 0.00479 | 3.14 | 4.08 | 122.36 | +1.97 |
| 160 | 3 | 0.18700 ± 0.00702 | 3.75 | 4.63 | 158.00 | -1.25 |
| 200 | 3 | 0.22260 ± 0.00997 | 4.48 | 5.33 | 195.02 | -2.49 |
| 240 | 3 | 0.25957 ± 0.00743 | 2.86 | 3.31 | 233.47 | -2.72 |
| 280 | 3 | 0.30820 ± 0.00672 | 2.18 | 2.46 | 284.04 | +1.44 |
| 320 | 3 | 0.34440 ± 0.00645 | 1.87 | 2.09 | 321.69 | +0.53 |
| 360 | 3 | 0.38650 ± 0.00436 | 1.13 | 1.24 | 365.48 | +1.52 |
| 400 | 3 | 0.42147 ± 0.00551 | 1.31 | 1.43 | 401.84 | +0.46 |
| 440 | 3 | 0.45410 ± 0.00364 | 0.80 | 0.87 | 435.78 | -0.96 |

Back-calculated concentrations use the 0-440 mg/dL calibration.

B Linear calibration

| Range (mg/dL) | n | Intercept | Slope | R² | Residual SD | Lack-of-fit p |
| --- | --- | --- | --- | --- | --- | --- |
| 0-440 | 39 | 0.03507921 | 0.0009615418 | 0.998398 | 0.005736 | 0.235957 |

Supplementary Table S3. Model performance across trajectory representations

Four trajectory representations are compared at a uniform threshold of 0.5000 and at their respective development OOF-selected thresholds. Each evaluation uses its own full-precision threshold. Thresholds and metrics are means across 100 repeats, with thresholds displayed to four decimal places. The accompanying Table_S3_model_metrics.tsv gives the minimum and maximum TP, FN, FP and TN counts across repeats. The summaries include all and drug-exposed observations; the accompanying files also provide no-drug controls, outer-fold results and confidence intervals. First and second differences are unchanged by centering. Confidence intervals and resampling are described in Supplementary Methods SM6.

A Development, all observations (N = 1,605). Growth (n = 961, 59.9%)

| Representation | Rule | Mean threshold | AUC | AP | Balanced accuracy |
| --- | --- | --- | --- | --- | --- |
| Original | Uniform | 0.5000 | 0.9945 | 0.9970 | 0.9736 |
| Original | OOF | 0.5192 | 0.9945 | 0.9970 | 0.9756 |
| T1-centered | Uniform | 0.5000 | 0.9968 | 0.9979 | 0.9732 |
| T1-centered | OOF | 0.4575 | 0.9968 | 0.9979 | 0.9755 |
| First difference | Uniform | 0.5000 | 0.9967 | 0.9980 | 0.9733 |
| First difference | OOF | 0.4316 | 0.9967 | 0.9980 | 0.9756 |
| Second difference | Uniform | 0.5000 | 0.9781 | 0.9859 | 0.9261 |
| Second difference | OOF | 0.5576 | 0.9781 | 0.9859 | 0.9298 |

B Development, drug-exposed observations (N = 1,400). Growth (n = 756, 54%)

| Representation | Rule | Mean threshold | AUC | AP | Balanced accuracy |
| --- | --- | --- | --- | --- | --- |
| Original | Uniform | 0.5000 | 0.9932 | 0.9955 | 0.9704 |
| Original | OOF | 0.5192 | 0.9932 | 0.9955 | 0.9725 |
| T1-centered | Uniform | 0.5000 | 0.9960 | 0.9967 | 0.9699 |
| T1-centered | OOF | 0.4575 | 0.9960 | 0.9967 | 0.9728 |
| First difference | Uniform | 0.5000 | 0.9959 | 0.9969 | 0.9699 |
| First difference | OOF | 0.4316 | 0.9959 | 0.9969 | 0.9731 |
| Second difference | Uniform | 0.5000 | 0.9725 | 0.9777 | 0.9171 |
| Second difference | OOF | 0.5576 | 0.9725 | 0.9777 | 0.9199 |

Supplementary Table S3 (continued)

C BALF, all wells (N = 144). Growth (n = 21, 14.6%)

| Representation | Rule | Mean threshold | AUC | AP | Balanced accuracy |
| --- | --- | --- | --- | --- | --- |
| Original | Uniform | 0.5000 | 0.9341 | 0.8172 | 0.5493 |
| Original | OOF | 0.5192 | 0.9341 | 0.8172 | 0.5482 |
| T1-centered | Uniform | 0.5000 | 0.9644 | 0.9024 | 0.8710 |
| T1-centered | OOF | 0.4575 | 0.9644 | 0.9024 | 0.8717 |
| First difference | Uniform | 0.5000 | 0.9453 | 0.8491 | 0.7993 |
| First difference | OOF | 0.4316 | 0.9453 | 0.8491 | 0.8087 |
| Second difference | Uniform | 0.5000 | 0.8061 | 0.6226 | 0.7892 |
| Second difference | OOF | 0.5576 | 0.8061 | 0.6226 | 0.7846 |

D BALF, drug-exposed wells (N = 120). Growth (n = 9, 7.5%)

| Representation | Rule | Mean threshold | AUC | AP | Balanced accuracy |
| --- | --- | --- | --- | --- | --- |
| Original | Uniform | 0.5000 | 0.8988 | 0.5827 | 0.5000 |
| Original | OOF | 0.5192 | 0.8988 | 0.5827 | 0.5014 |
| T1-centered | Uniform | 0.5000 | 0.9361 | 0.7112 | 0.7219 |
| T1-centered | OOF | 0.4575 | 0.9361 | 0.7112 | 0.7292 |
| First difference | Uniform | 0.5000 | 0.8905 | 0.5703 | 0.6598 |
| First difference | OOF | 0.4316 | 0.8905 | 0.5703 | 0.6590 |
| Second difference | Uniform | 0.5000 | 0.6335 | 0.2471 | 0.6650 |
| Second difference | OOF | 0.5576 | 0.6335 | 0.2471 | 0.6515 |

Uniform denotes the shared threshold of 0.5000; OOF denotes the threshold selected from pooled development out of fold scores for each representation. The same development-set OOF predictions were used for both threshold selection and performance evaluation. ROC-AUC and AP are calculated from the same continuous scores under both rules. In growth-only subsets, ROC-AUC, balanced accuracy, specificity and false-positive rate are undefined (NA). Outer folds have overlapping training sets; BALF wells are technical replicates within eight specimens from six patients. N, number of wells; growth, number of wells labeled as growth; AUC, area under the receiver operating characteristic curve; AP, average precision. BALF, bronchoalveolar lavage fluid; T1, first reading. TP, true positive; FN, false negative; FP, false positive; TN, true negative. Growth assessed manually after overnight incubation is the positive reference class.

Supplementary Table S4. Predicted and reference MICs for cultured isolates

The complete 228-row table is provided as the supplementary TSV attachment tables/Table_S4_reference_MIC.tsv.

Predicted and reference MICs are reported for 228 isolate-drug combinations (Methods 2.6 and 2.13). For each combination, the table shows the MIC predicted most often among repeats that yielded a result. All 100 repeats yielded a MIC except for ATCC25922 with AMP_SUL or AZT (99 repeats each). Frequencies for individual MIC values are provided in Table_S4_reference_MIC.tsv. Combination-drug concentrations are listed in drug-name order, with AMP:SUL at 2:1. MIC agreement was assessed using the ampicillin component for AMP_SUL and the levofloxacin component for AMK_LEV (Methods 2.14). Agreement categories are EX (exact agreement), EA (within one twofold dilution, excluding EX) and OEA (outside essential agreement); overall EA includes exact matches. NIT series used shared no-drug controls; the ATCC25922-AZT series used the same-day, same-isolate AMP_SUL control. An asterisk (*) marks combinations with skipped concentrations in at least one repeat (n = 27, 11.8%). MIC, minimum inhibitory concentration; d, log₂(predicted/reference MIC).

Supplementary Table S5. Comparison of susceptibility-related approaches

Supplementary Table S5. Selected molecular and phenotypic approaches compared by output, starting material, assay interval, evaluation design and equipment. The cited studies and product descriptions use different samples and endpoints; times refer to the intervals specified in each row. Clinical turnaround also includes preceding and subsequent workflow steps. The Category column identifies nucleic acid detection, conventional and commercial phenotypic AST, investigational rapid phenotypic methods, and GT-AST. Commercial systems may also provide rapid testing. GT-AST resource use is detailed in Table S6. AST, antimicrobial susceptibility testing; BMD, broth microdilution; MIC, minimum inhibitory concentration; GT-AST, glucose-turbidity antimicrobial susceptibility testing; BALF, bronchoalveolar lavage fluid; BCID2, BIOFIRE Blood Culture Identification 2 Panel; PCR, polymerase chain reaction; IR-ATR, infrared attenuated total reflection; PCA, principal component analysis; FTIR, Fourier-transform infrared spectroscopy; ATR, attenuated total reflection; LC-MS, liquid chromatography-mass spectrometry; UHPLC-MS, ultra-high-performance liquid chromatography-mass spectrometry; CFU, colony-forming units.

| Category | Approach / source | Measurement and output | Sample preparation and timing | Study unit and evaluation | Equipment and operations | Key features |
| --- | --- | --- | --- | --- | --- | --- |
| Nucleic acid detection | BIOFIRE BCID2 (FDA, 2020) | Multiplex PCR identification of selected organisms and resistance genes. | Positive blood culture; approximately 1 h of testing after culture positivity. | 1,074 prospective positive blood-culture specimens, supplemented by archived and seeded specimens; analyte-level agreement. | FILMARRAY 2.0 or TORCH, single-use pouches and automated processing after sample/buffer loading. | Provides organism and resistance-marker information complementary to phenotypic measurements. |
| Conventional and commercial phenotypic AST | Conventional BMD (EUCAST, 2024; 2026, BMD recommendations) | MIC from visible growth inhibition across antibiotic concentrations. | Cultured isolates; standardized inoculum and overnight incubation. Isolation precedes testing. | EUCAST reference-method guidance. | Microdilution plates, incubator and manual reading, with purity, inoculum and growth-control checks. | Conventional growth-inhibition framework and principal incubation-time comparator. |
|  | VITEK 2 (FDA K183415, 2019) | Automated growth monitoring in AST cards; MIC and susceptibility category. | Standardized isolate suspension; up to 18 h of instrument incubation for the cited Gram-negative imipenem card, after isolation and inoculum preparation. | Regulatory evaluation of the specified Gram-negative imipenem card. | VITEK 2 instrument, disposable AST cards and standardized inoculum preparation. | Automated phenotypic AST from cultured isolates. |
|  | VITEK REVEAL (FDA K230675, 2024; Gitman et al., 2026) | Phenotypic AST with MIC reporting for Gram-negative bloodstream infections. | Positive blood culture; mean time to result from AST test initiation: 6.0 h for sequential results and 7.9 h for the final result. | Six-centre evaluation of 553 Gram-negative positive blood cultures; comparison with MicroScan. | Dedicated automated instrument and test consumables. | Rapid commercial phenotypic AST starting from positive blood cultures. |
| Investigational rapid phenotypic methods | Sturm et al., 2024 | Cantilever nanomotion and classification models; susceptibility calls at selected drug concentrations. | Spiked or patient positive blood cultures, followed by sample processing and cell attachment. Recording: 4 h at room temperature; 2 h at 37 °C for ceftazidime-avibactam. | Development: 364 isolates. Independent testing: 223 strains, including evaluation in a clinical setting; accuracy 89.5-98.9%. | Cantilever platform, cell-attachment reagents and signal processing. | Nanomotion-based classification evaluated on independent strains. |
|  | Jafari et al., 2024 | IR-ATR glucose monitoring with PCA-based discrimination of antibiotic responses. | Overnight culture and washing, followed by exposure at 10^8^CFU/mL; glucose changes assessed after 2 h. | Four strains and four antibiotics; proof-of-concept experiments. | FTIR spectrometer with ATR accessory; serial sampling and spectral analysis. | Glucose-based response measurement; GT-AST pairs glucose with turbidity and later growth. |
|  | Rydzak et al., 2022 | LC-MS susceptibility calls based on inhibition of glucose consumption (*S. aureus*/*Enterococcus*), succinate production (*E. coli*/*K. pneumoniae*) or nicotinate production (*S. pneumoniae*). | Prepared isolates: 4 h incubation, sample processing and 5-min chromatography per sample. A spiked positive-blood-culture workflow was also examined. | AST: 246 isolates (80 training, 166 test); comparison with VITEK 2. | UHPLC-MS, with centrifugation, supernatant extraction and sequential chromatography. | Metabolic AST with species-dependent markers and a separate test set. |
|  | Riester et al., 2024 | Resazurin differential pulse voltammetry combined with impedance for growth and susceptibility calls. | Diluted spiked serum; combined-method growth detection at 4.94 ± 0.28 to 10.22 ± 1.59 h. | Seven strains, two antibiotics (kanamycin and oxytetracycline), triplicate tests (42 total). Per-replicate sensitivity 94.44% and specificity 95.83% against agar diffusion. | 3D-printed chambers, screen-printed electrodes, potentiostat and computational analysis. | Combined readouts tested in experimentally spiked serum. |
| GT-AST | GT-AST (this study) | Paired residual-glucose and turbidity measurements; growth prediction and concentration-series MIC estimates. | Cultured isolates; BALF tested without prior isolation. Eight time points at 30-135 min after inoculation; each glucose readout also requires sample processing and color development. Overnight growth readings provide reference outcomes. Sample-to-report time remains to be evaluated. | Isolate-grouped evaluation (43 isolates, 228 isolate-drug pairs). Mean exact agreement 64.8%, EA 87.1%. Retrospective BALF evaluation (8 specimens, 6 patients). For the four culture-positive specimens, mean exact agreement and EA were both 99.0%. BALF species inputs came from subsequently cultured isolates. | Conventional microplate reader, incubator, centrifuge and heating equipment; manual sampling and color development. | Pairs two early response readouts and evaluates their relation to later growth inhibition. |

Supplementary Table S6. Reagent costs and equipment requirements for the manual BALF workflow

Estimated reagent cost per BALF specimen

| Reagent | Quantity per specimen | Reference price (USD / package) | Cost per specimen (USD) |
| --- | --- | --- | --- |
| Glacial acetic acid | 27-33.75 mL | 26.95 / 5,000 mL | 0.1455-0.1819 |
| o-Toluidine | 1.8-2.25 mL | 8.56 / 500 mL | 0.0308-0.0385 |
| Thiourea | 0.045-0.05625 g | 7.52 / 500 g | 0.0007-0.0008 |
| Boric acid | 0.072-0.09 g | 23.82 / 500 g | 0.0034-0.0043 |
| Beef powder | 0.12-0.15 g | 48.24 / 500 g | 0.0116-0.0145 |
| Casein hydrolysate | 1.05-1.3125 g | 134.00 / 500 g | 0.2814-0.3518 |
| Levofloxacin | 0.48-0.6 mg | 49.13 / 5 g | 0.0047-0.0059 |
| D-(+)-Glucose | 0.18-0.225 g | 7.74 / 250 g | 0.0056-0.0070 |
| Approximate total | — | — | 0.50-0.60 |

One test denotes one BALF specimen evaluated across seven LEV conditions (0, 0.5, 1, 2, 4, 6 and 8 µg/mL), with three technical replicates and eight paired glucose-turbidity readings. Four specimens share two fully utilized 96-well culture plates; shared controls and preparation allowances are included in the per-specimen allocation. BALF, bronchoalveolar lavage fluid; LEV, levofloxacin.

Required equipment comprises a microplate reader, a shaking incubator, a plate centrifuge, a thermocycler and a multichannel pipette.

Reference equipment prices

| Equipment | Pricing unit | Reference price (USD) |
| --- | --- | --- |
| Microplate reader (Single fixed wavelength) | 1 unit | 1,489 |
| Shaking incubator | 1 unit | 1,191 |
| Plate centrifuge with plate rotor | 1 set | 596 |
| Thermocycler | 1 unit | 1,191 |
| Multichannel pipette | 1 pipette | 1,042 |

Supplementary Table S7. Comparison of BALF experimental findings, predicted MICs and hospital results

| Specimen | Plating result | No-drug control wells with growth, n (%) | Hospital detection (organism) | Hospital MIC (µg/mL) | Most frequent predicted MIC (µg/mL) | MIC difference (twofold dilutions) |
| --- | --- | --- | --- | --- | --- | --- |
| BALF001 | P | 3 (100) | P (*Stenotrophomonas maltophilia*) | ≤2 (S) | ≤0.5 (S) | 0 |
| BALF002 | N | 0 (0) | N | — | ≤0.5 | — |
| BALF003 | P | 3 (100) | P (*Enterococcus faecalis*) | ≤2 (S) | 1 (S) | 0 |
| BALF004 | P | 3 (100) | P (*Pseudomonas aeruginosa*) | ≤1 (S) | ≤0.5 (S) | 0 |
| BALF005 | N | 0 (0) | N | — | — | — |
| BALF006 | P | 3 (100) | P (*Pseudomonas aeruginosa*) | ≤1 (S) | ≤0.5 (S) | 0 |
| BALF007 | N | 0 (0) | N | — | — | — |
| BALF008 | N | 0 (0) | N | — | — | — |

P, positive; N, negative. No-drug control growth is the number and percentage of the three control wells that were turbid after overnight incubation. All MICs are for levofloxacin (LEV). Parentheses after MICs indicate susceptibility categories: S, susceptible. Hospital categories are as reported; predicted MIC categories are based on the species-specific numerical breakpoints listed in CLSI M100, 36th edition (CLSI, 2026). A dash (—) indicates that no MIC was reported or no MIC comparison was applicable.

Predicted MICs used the full model at a growth-score threshold of 0.5 and the three-replicate majority-vote procedure in Methods 2.13. The table shows the MIC predicted most often among repeats that yielded a result. Individual predictions from all 100 repeats are provided in Table_S7_repeat_MIC.tsv. The hospital reported a levofloxacin MIC of ≤2 µg/mL for BALF001. The full model correctly classified all 18 wells of BALF004 in every repeat. In BALF001, one to three of the three growth wells were missed per repeat at 0.5 µg/mL. At 1 µg/mL, all three were missed in 99 of the 100 repeats and two were missed in the remaining repeat. In BALF003, the 100 majority calls at 1 µg/mL were no growth (n = 85, 85%) or growth (n = 15, 15%), yielding predicted MICs of 1 and 2 µg/mL, respectively.

MIC difference is the predicted minus hospital result in twofold dilutions, calculated using the CLSI comparison rules in Methods 2.14. Exact agreement and EA follow the definitions in that section.

Eight specimens came from six patients. The specimens with no growth in the no-drug controls (n = 4, 50%) had no pathogen detected by the hospital. These specimens required no MIC comparison. Among them, only BALF002 yielded a predicted MIC, in the 10 of 100 repeats in which the control was predicted to grow. All four culture-positive specimens from three patients had hospital MICs. BALF006 yielded a predicted MIC of ≤0.5 µg/mL in 96 evaluations. In the other four, the model predicted no growth in its no-drug control and yielded no MIC. Across 100 evaluations, mean MIC prediction coverage for the four culture-positive specimens was 99.0%. Exact agreement and EA with hospital MICs also averaged 99.0%. Plating and hospital bacterial detection agreed for all eight specimens. LEV classifications based on observed growth agreed with hospital categories for all four culture-positive specimens. BALF001 and BALF004 were collected during ICU care and the remaining specimens during follow-up after treatment in a general ward. BALF001/BALF005 and BALF004/BALF006 were sequential specimens from two patients. BALF, bronchoalveolar lavage fluid; MIC, minimum inhibitory concentration; ICU, intensive care unit.

Supplementary Table S8. Growth prediction and MIC results across input configurations

Blocks A and B summarize drug-exposed wells from cultured isolates and BALF, respectively; block C summarizes MIC results. The development cohort comprised 43 isolates, and BALF evaluation included eight specimens from six patients. Model inputs and implementation are described in Supplementary Methods SM4.

Glucose and Turbidity denote the corresponding trajectories; S, species category; D, antibiotic components; C, concentration level. Full model: Glucose + Turbidity + S + D + C.

The six supplementary TSV attachments for the interval estimates and paired comparisons are provided in tables/: Table_S8_development_intervals.tsv, Table_S8_development_paired_intervals.tsv, Table_S8_BALF_intervals.tsv, Table_S8_BALF_paired_intervals.tsv, Table_S8_MIC_intervals.tsv and Table_S8_MIC_paired_intervals.tsv.

A. Cultured-isolate drug-exposed wells (N = 1,400). Growth (n = 756, 54%), no growth (n = 644, 46%)

| Inputs | ROC-AUC | AP | Accuracy (%) | Balanced accuracy (%) | Growth recall (%) | False-positive rate (%) |
| --- | --- | --- | --- | --- | --- | --- |
| Full model | 0.99596 | 0.99671 | 96.97 | 96.99 | 96.67 | 2.68 |
| Glucose + Turbidity + D + C | 0.99529 | 0.99618 | 97.01 | 97.06 | 96.52 | 2.41 |
| Turbidity + S + D + C | 0.99434 | 0.99557 | 96.26 | 96.33 | 95.56 | 2.91 |
| Glucose + S + D + C | 0.98442 | 0.98642 | 94.40 | 94.40 | 94.45 | 5.65 |
| S + D + C | 0.83990 | 0.84130 | 74.03 | 74.33 | 70.58 | 21.91 |
| Glucose + Turbidity | 0.99585 | 0.99673 | 97.07 | 97.12 | 96.48 | 2.24 |
| Glucose + Turbidity + S + C | 0.99611 | 0.99687 | 97.04 | 97.10 | 96.34 | 2.15 |
| Glucose + Turbidity + S + D | 0.99595 | 0.99672 | 96.93 | 96.98 | 96.36 | 2.41 |

Supplementary Table S8 (continued)

B. BALF drug-exposed wells (N = 120). Growth (n = 9, 7.5%), no growth (n = 111, 92.5%)

| Inputs | ROC-AUC | AP | Accuracy (%) | Balanced accuracy (%) | Growth recall (%) | False-positive rate (%) |
| --- | --- | --- | --- | --- | --- | --- |
| Full model | 0.93606 | 0.71118 | 94.92 | 72.19 | 45.44 | 1.07 |
| Glucose + Turbidity + D + C | 0.96506 | 0.80612 | 95.69 | 73.42 | 47.22 | 0.38 |
| Turbidity + S + D + C | 0.97369 | 0.86421 | 95.22 | 68.17 | 36.33 | 0.00 |
| Glucose + S + D + C | 0.77069 | 0.43531 | 85.88 | 61.79 | 33.44 | 9.87 |
| S + D + C | 0.73613 | 0.16759 | 75.25 | 55.53 | 32.33 | 21.27 |
| Glucose + Turbidity | 0.95857 | 0.79117 | 95.78 | 73.57 | 47.44 | 0.31 |
| Glucose + Turbidity + S + C | 0.95078 | 0.70649 | 93.93 | 74.11 | 50.78 | 2.57 |
| Glucose + Turbidity + S + D | 0.92352 | 0.72015 | 95.63 | 74.16 | 48.89 | 0.58 |

Growth is the positive reference class. ROC-AUC and average precision (AP) use continuous scores; classification metrics use a threshold of 0.5. Growth recall is TP/(TP + FN), and the false-positive rate is FP/(TN + FP). Balanced accuracy averages growth recall and specificity. Metrics are means across 100 evaluations. The panel headings give the numbers of observed growth and no-growth wells used to calculate growth recall and the false-positive rate, respectively. Patient-cluster 95% intervals and paired differences against the full model are provided in Table_S8_BALF_intervals.tsv and Table_S8_BALF_paired_intervals.tsv, respectively; sample-cluster sensitivity intervals are also retained. Corresponding cultured-isolate intervals and paired differences are provided in Table_S8_development_intervals.tsv and Table_S8_development_paired_intervals.tsv. Confidence intervals and paired comparisons are described in Supplementary Methods SM6.

Supplementary Table S8 (continued)

C. MIC agreement

C1. Cultured-isolate MIC agreement

| Inputs (mean MIC prediction coverage, %) | Mean EX (%) | Mean EA (%) |
| --- | --- | --- |
| Full model (99.99%) | 64.78 | 87.05 |
| Glucose + Turbidity + D + C (100.00%) | 64.39 | 86.35 |
| Turbidity + S + D + C (100.00%) | 64.44 | 86.32 |
| Glucose + S + D + C (99.97%) | 56.88 | 83.10 |
| S + D + C (100.00%) | 29.32 | 55.40 |
| Glucose + Turbidity (99.91%) | 65.09 | 87.38 |
| Glucose + Turbidity + S + C (100.00%) | 64.45 | 88.03 |
| Glucose + Turbidity + S + D (99.86%) | 64.86 | 87.38 |

C2. BALF MIC agreement

| Inputs (mean MIC prediction coverage, %) | Mean EX (%) | Mean EA (%) |
| --- | --- | --- |
| Full model (99.00%) | 99.00 | 99.00 |
| Glucose + Turbidity + D + C (80.25%) | 80.25 | 80.25 |
| Turbidity + S + D + C (78.25%) | 78.25 | 78.25 |
| Glucose + S + D + C (100.00%) | 88.75 | 97.75 |
| S + D + C (100.00%) | 50.00 | 50.00 |
| Glucose + Turbidity (75.00%) | 75.00 | 75.00 |
| Glucose + Turbidity + S + C (99.50%) | 99.50 | 99.50 |
| Glucose + Turbidity + S + D (75.00%) | 75.00 | 75.00 |

Supplementary Table S8 (continued)

C3. Paired comparisons across 228 isolate-drug combinations

| Inputs | Reference MICs (N) | Full model EX (%) | Configuration EX (%) | Full model EA (%) | Configuration EA (%) |
| --- | --- | --- | --- | --- | --- |
| Glucose + Turbidity + D + C | 228 | 64.78 | 64.39 | 87.05 | 86.35 |
| Turbidity + S + D + C | 228 | 64.78 | 64.44 | 87.05 | 86.32 |
| Glucose + S + D + C | 228 | 64.78 | 56.88 | 87.05 | 83.10 |
| S + D + C | 228 | 64.78 | 29.32 | 87.05 | 55.40 |
| Glucose + Turbidity | 228 | 64.78 | 65.09 | 87.05 | 87.38 |
| Glucose + Turbidity + S + C | 228 | 64.78 | 64.45 | 87.05 | 88.03 |
| Glucose + Turbidity + S + D | 228 | 64.78 | 64.86 | 87.05 | 87.38 |

C4. Paired comparisons across four culture-positive BALF specimens

| Inputs | Reference MICs (N) | Full model EX (%) | Configuration EX (%) | Full model EA (%) | Configuration EA (%) |
| --- | --- | --- | --- | --- | --- |
| Glucose + Turbidity + D + C | 4 | 99.00 | 80.25 | 99.00 | 80.25 |
| Turbidity + S + D + C | 4 | 99.00 | 78.25 | 99.00 | 78.25 |
| Glucose + S + D + C | 4 | 99.00 | 88.75 | 99.00 | 97.75 |
| S + D + C | 4 | 99.00 | 50.00 | 99.00 | 50.00 |
| Glucose + Turbidity | 4 | 99.00 | 75.00 | 99.00 | 75.00 |
| Glucose + Turbidity + S + C | 4 | 99.00 | 99.50 | 99.00 | 99.50 |
| Glucose + Turbidity + S + D | 4 | 99.00 | 75.00 | 99.00 | 75.00 |

C1 and C2 show the mean percentage of isolate-drug combinations or culture-positive BALF specimens with a predicted MIC for each input configuration. Prediction coverage, exact agreement (EX) and essential agreement (EA) were calculated in each evaluation using all 228 combinations or all four specimens, then averaged across the 100 evaluations. A missing prediction counted as a failure. C3 and C4 compare each configuration with the full model on the same combinations or specimens. Supplementary Table S7 gives the full-model BALF results. Table_S8_MIC_intervals.tsv and Table_S8_MIC_paired_intervals.tsv give confidence intervals for each configuration and for its difference from the full model (Methods 2.13-2.14 and Supplementary Methods SM6).

In cultured-isolate drug-exposed wells, mean balanced accuracy was 96.99% for the full model and 74.33% for species, drug and concentration inputs alone. In drug-exposed BALF wells, the full model and the model without species input had mean accuracies of 94.92% and 95.69%, respectively. Of nine observed growth wells, the full model detected three to six per repeat and the model without species input detected three to five. Of 111 no-growth wells, the models misclassified zero to three and zero to two per repeat, respectively. This comparison evaluates the effect of including species input when six BALF categories were absent from development. These exploratory estimates describe input-related differences in growth detection, false-positive calls and MIC estimation within the current cohorts. Paired curves characterize distinct glucose and turbidity responses during drug exposure; their phenotypic information and prediction performance provide complementary perspectives. BALF, bronchoalveolar lavage fluid; MIC, minimum inhibitory concentration; T1, first reading; ROC-AUC, area under the receiver operating characteristic curve; EA, essential agreement. TP, true positive; FN, false negative; FP, false positive; TN, true negative. Growth assessed manually after overnight incubation is the positive reference class.

Supplementary Table S9. Concentration categories and actual concentrations

Concentration level

| Drug condition | 0 | 1 | 2 | 3 | 4 | 5 | 6 | 7 | 8 | 9 |
| --- | --- | --- | --- | --- | --- | --- | --- | --- | --- | --- |
| AMK | 0 | 2 | - | 4 | - | 8 | - | 16 | - | 32 |
| AMK + LEV | 0 | 1 | - | - | - | 4 | - | - | - | 32 |
| AMP + SUL | 0 | 1 | 2 | - | 4 | 8 | 16 | - | 32 | 64 |
| AZT | 0 | 1 | 2 | - | 4 | 8 | 16 | - | 32 | 64 |
| CFZ | 0 | 1 | 2 | - | 4 | 8 | 16 | - | 32 | 64 |
| CIP | 0 | 0.25 | - | 0.5 | - | 1 | - | 2 | - | 4 |
| CRO | 0 | 1 | 2 | - | 4 | 8 | 16 | - | 32 | 64 |
| ERY | 0 | 0.25 | - | 0.5 | 1 | - | 2 | 4 | - | 8 |
| LEV | 0 | 0.5 | - | 1 | 2 | - | 4 | 6 | - | 8 |
| NIT | 0 | 2 | 4 | 8 | 16 | - | 32 | 64 | 128 | 256 |

Values are antibiotic concentrations in µg/mL for each complete drug condition. Level 0 denotes no drug; a dash denotes an unused category for that condition. Levels 1-9 identify the ordered concentration categories and do not encode dose intervals. For example, level 3 corresponds to 4 µg/mL AMK, 0.5 µg/mL CIP and 1 µg/mL LEV. For BALF analyses, level 7 (6 µg/mL LEV) was omitted and the other level identifiers were retained. MIC derivation and dilution-step comparisons used the actual concentrations. AMK, amikacin; AMK + LEV, amikacin plus levofloxacin; AMP + SUL, ampicillin plus sulbactam; AZT, aztreonam; CFZ, cefazolin; CIP, ciprofloxacin; CRO, ceftriaxone; ERY, erythromycin; LEV, levofloxacin; NIT, nitrofurantoin. BALF, bronchoalveolar lavage fluid; MIC, minimum inhibitory concentration.

Supplementary Text

Methodology

SM1. IPTG induction experiments

We examined glucose consumption across isopropyl β-D-1-thiogalactopyranoside (IPTG) concentrations under antibiotic exposure using engineered *Escherichia coli* 10β carrying pET-15b-KanR and *Bacillus subtilis* BS168 carrying pHT43-ErmR. Kanamycin- and erythromycin-resistance genes from pJOE8999 and pAX01, respectively, were inserted into the corresponding expression vectors. Recombinant constructs were checked by colony PCR and sequencing, and strains carrying the corresponding empty vectors served as controls.

*B. subtilis* was precultured to logarithmic phase, whereas *E. coli* was precultured overnight in liquid medium containing the IPTG concentration used in its subsequent exposure condition. Before inoculation, culture density was estimated from optical density at 600 nm measured with a SmartSpec Plus spectrophotometer (Bio-Rad, USA). For *E. coli*, the conversion setting stored in the instrument was used. Cultures were then inoculated into Luria-Bertani (LB) medium at a target final inoculum of approximately 5 × 10⁵ colony-forming units (CFU)/mL.

The exposure medium contained 10 µg/mL kanamycin for *E. coli* or 50 µg/mL erythromycin for *B. subtilis*. Recombinant strains were tested at IPTG concentrations of 0, 0.01, 0.05, 0.1, 0.5, 1, 2.5, 5 and 10 mM; the displayed empty-vector controls received no IPTG. Glucose concentrations were measured with Accu-Chek Active blood glucose test strips (Roche) and recorded manually. The time courses in Supplementary Fig. S1A and B used readings at 0 h and every 2 h from 4 to 8 h for *E. coli*, and at 0 h and every 2 h from 10 to 18 h for *B. subtilis*.

For each bacterial model, the displayed glucose concentrations are condition means from a single experiment. Average glucose consumption rates were calculated for each strain and IPTG concentration from the mean time course, using the decrease from the initial concentration to the last strictly positive reading divided by the elapsed time between those readings (mmol/(L·h)). Terminal zero readings were excluded from this calculation. For *E. coli* and *B. subtilis* separately, rates at positive IPTG concentrations were fitted by unweighted nonlinear least squares to a four-parameter Hill model with freely estimated Bottom, Top, EC_50_ and Hill slope (Supplementary Fig. S1C and D, respectively). The 0 mM condition was excluded from fitting. Bottom and Top are the fitted lower and upper plateaus, respectively; EC_50_ is the IPTG concentration at the midpoint between these plateaus, in mM, and Hill slope describes the steepness of the fitted response. The source data and analysis code accompany the supplementary material.

SM2. Glucose colorimetry

The 6% o-toluidine reagent was prepared by dissolving 1.5 g thiourea in 500 mL glacial acetic acid, adding 60 mL o-toluidine and 40 mL saturated boric acid solution, and bringing the volume to 1 L with glacial acetic acid. The reagent was stored at 4 °C protected from light in a container sealed with laboratory sealing film. Saturated boric acid solution was prepared by heating 60 g boric acid in 1 L water, allowing the solution to stand overnight and filtering the supernatant.

Glucose color development was compared across heating conditions and sample:reagent volumes using 400 mg/dL glucose in Mueller-Hinton (MH) medium. Heating temperatures of 92, 95 and 98 °C, heating times of 10, 12 and 14 min, and sample:o-toluidine reagent volumes of 20:170, 40:150 and 60:130 µL were combined in a 3 × 3 × 3 design. For each of the 27 conditions, the same source solution was dispensed into eight separate wells and heated within the same batch, providing eight technical replicates. Absorbance was measured at 620 nm using an infinite F50 microplate reader. The selected conditions were 95 °C for 12 min with 40 µL sample and 150 µL reagent (Supplementary Fig. S2 and Supplementary Table S1).

Under the selected conditions, glucose standards prepared in MH medium were measured over 0-440 mg/dL in 40 mg/dL increments. Concentrations refer to the MH solution before mixing with reagent. The zero-concentration MH standard had six technical replicate wells, and each nonzero concentration had three. The 480 mg/dL level was excluded because one of its three planned wells was unfilled. Water-control readings were recorded separately from the zero-concentration MH standards.

A single linear calibration curve was fitted to all 39 individual absorbance readings over 0-440 mg/dL by ordinary least squares, with a freely estimated intercept and equal weight for each well. Residuals were calculated as observed minus fitted absorbance. Linear lack of fit was assessed against the pure error estimated from replicate wells at each concentration. Glucose concentrations were back-calculated as (absorbance − intercept)/slope. For each nonzero standard concentration, relative deviation was calculated as 100 × (mean back-calculated concentration − nominal concentration)/nominal concentration.

Absorbance for each condition or concentration was summarized as the arithmetic mean and sample standard deviation (SD). The coefficient of variation (CV) was calculated as 100 × SD/mean. For nonzero glucose standards, net-signal CV used the same SD and the mean absorbance after subtraction of the mean of the six zero-concentration MH wells (Supplementary Fig. S3 and Supplementary Table S2).

SM3. Ordination features

Principal coordinates analysis included the observations in the response map and quality-eligible combination-drug observations. Each observation was represented by 22 features: the seven glucose decreases from the first glucose reading and seven turbidity increases from the first turbidity reading at sampling points 2-8; the sum of these changes for each channel; the first-to-last change for each channel; each channel's maximum-minus-minimum range; and its least-squares slope against sampling index. These sums summarized changes across sampled time points. For the series-level analysis, observations were grouped by isolate and complete drug context. The mean, maximum, minimum and population standard deviation of each of the 22 features were calculated across tested concentrations, and the number of concentrations was appended, giving 89 features for each series.

SM4. Computational implementation

Model architecture and training

Each input block was encoded as a 16-dimensional vector. Each trajectory encoder used two one-dimensional convolutional layers with 16 channels, kernel width 3, padding 1 and Gaussian error linear unit (GELU) activations. Global mean and maximum pooling were concatenated and passed through two fully connected layers of width 16, with GELU activation and dropout of 0.1 between them. Each indicator block used a separate two-layer fully connected encoder of width 16 with the same activation and dropout. The five vectors were combined by two-head attention using a learned query, linear key and value projections, and softmax weights over the input blocks. The resulting 16-dimensional vector passed through a prediction head with 64 hidden units, GELU activation, dropout of 0.2 and one output logit. A sigmoid transformation gave the model growth score.

Models were trained using AdamW (Loshchilov and Hutter, 2019) with a learning rate of 0.002, weight decay of 0.0001, batch size 64 and a maximum of 150 epochs. Binary cross-entropy loss weighted the positive class by the ratio of negative to positive observations in the optimization partition. Gradient norms were clipped at 5. For early stopping, each model used a seven-fold StratifiedGroupKFold split with shuffling and its own seed. The first split with both outcome classes in the optimization and stopping subsets was selected. Training stopped after 25 epochs without an improvement in stopping-set area under the receiver operating characteristic curve (ROC-AUC) greater than 10⁻⁵, and the weights at the best stopping epoch were retained.

Trajectory representations and input configurations

Subtracting the first reading (T1) leaves first and second differences unchanged, so only the original trajectories, T1-centered trajectories and the two difference orders were fitted. The summary curves and per-well displays in Supplementary Figs. S7-S9 each show one result per distinct input. Classification results in these figures use thresholds selected from pooled development out of fold (OOF) scores in each repeat. The mean thresholds were 0.5192 for original trajectories, 0.4575 for T1-centered trajectories, 0.4316 for first differences and 0.5576 for second differences. Supplementary Table S3 reports these results alongside those at the uniform threshold of 0.5000. Thresholds are displayed to four decimal places; calculations use the saved full-precision values.

Within each repeat, eight input configurations were evaluated using the same isolate-grouped partitions, five random seeds and training settings. Each model retained encoders for its listed inputs. Trajectories comprised eight T1-centered readings at 30-135 min.

Software environment

Modeling used Python 3.12.3, NumPy 2.3.5, pandas 2.3.3, scikit-learn 1.6.1 and PyTorch 2.8.0. Training was performed on the CPU with two PyTorch threads and deterministic algorithms enabled.

SM5. MIC comparison

MIC estimation used the full model's isolate-grouped out of fold predictions for 43 cultured isolates, including two ATCC reference strains. Scores ≥0.5 predicted growth, and MICs were derived as described in Methods 2.13. Each prediction was matched to the reference MIC for the same isolate-drug combination (Methods 2.6). Predicted and reference MICs are listed in Supplementary Table S4.

MIC differences for cultured isolates were expressed as d = log₂(p/r), where p and r denote predicted and reference MICs, respectively. For off-scale MICs, the reported concentration limit was used. Exact agreement, EA and OEA corresponded to d = 0, |d| ≤ 1 and |d| > 1, respectively. BALF MIC agreement was assessed using the CLSI Breakpoint Implementation Toolkit (CLSI, 2023), which incorporates off-scale MICs in the calculation of dilution differences and EA.

SM6. Confidence intervals and paired comparisons

For the full-model cultured-isolate MIC agreement analysis, confidence intervals used 20,000 isolate-level bootstrap replicates generated with NumPy PCG64. Each replicate sampled 43 isolates with replacement and included all their combinations; descriptive subsets used their own isolate sets. Agreement was calculated separately for each training repeat and averaged within each bootstrap draw. The 2.5th and 97.5th percentiles were calculated by linear interpolation. Predictions remained fixed, and all 20,000 replicates had valid denominators.

For the trajectory-representation comparisons in Supplementary Table S3, confidence intervals used 20,000 shared cluster bootstrap draws, resampling isolates for development and patients for BALF, with specimen resampling as a BALF sensitivity analysis. Models and thresholds remained fixed during resampling.

For the input-configuration comparisons in Supplementary Table S8, patient-cluster intervals were used for BALF classification, with sample-cluster intervals retained as a sensitivity analysis. The MIC and classification comparisons used the same 20,000 cluster draws (NumPy PCG64); percentile limits were calculated by linear interpolation from defined replicates. The files report the applicable denominators and valid and undefined replicate counts. Paired differences are the listed configuration minus the full model.

Results

SR1. IPTG induction experiments

To assess glucose consumption as a readout of bacterial responses under antibiotic exposure, we compared engineered bacteria across IPTG concentrations at a fixed antibiotic concentration for each model. In both *E. coli* and *B. subtilis*, higher IPTG concentrations were generally associated with greater glucose depletion, whereas the empty-vector controls and uninduced recombinant cultures showed comparatively small changes over the displayed time courses (Supplementary Fig. S1A and B). Average glucose consumption rates derived from these time courses also generally increased with IPTG concentration (Supplementary Fig. S1C and D). These results provide initial support for glucose consumption as a readout of metabolic responses in these engineered bacteria under antibiotic exposure.

SR2. Glucose colorimetry

At 400 mg/dL glucose, extending heating at 95 °C from 10 to 12 min increased mean absorbance by 19.33-22.52% across the three sample:reagent volume combinations (Supplementary Fig. S2 and Supplementary Table S1). At 98 °C, extending heating from 12 to 14 min decreased mean absorbance by 4.91-8.43%. The CVs among the eight technical replicate wells for each of the 27 conditions ranged from 0.92% to 4.27%. For the selected conditions of 95 °C, 12 min and 40:150 µL, absorbance was 0.41156 ± 0.00871 (mean ± SD; n = 8; CV = 2.12%).

Under the selected conditions, absorbance at 620 nm increased approximately linearly with glucose concentration over 0-440 mg/dL (Supplementary Fig. S3A and B and Supplementary Table S2). The calibration equation was A_620_ = 0.03507921 + 0.0009615418 × C, where C is glucose concentration in mg/dL (R² = 0.998398; lack-of-fit p = 0.2360). The maximum absolute relative deviation of the mean back-calculated concentration from the nominal concentration was 3.01% over 40-440 mg/dL. Within this nonzero concentration range, CVs were 0.80-4.48% for raw absorbance and 0.87-5.88% for the net signal after subtraction of the zero-concentration MH mean (Supplementary Fig. S3C).

SR3. MIC agreement

All 228 isolate-drug combinations from 43 cultured isolates had reference MICs. Across 100 evaluations, the model provided MIC estimates for an average of 99.99% of these combinations. Mean exact agreement was 64.8% (isolate-bootstrap 95% confidence interval, 58.8-70.9%) and mean EA was 87.1% (82.0-91.6%). Descriptive analyses excluding the two ATCC strains or retaining only single-agent treatments gave mean EA of 87.9% (216 combinations) and 87.4% (196 combinations), respectively. Supplementary Table S4 lists all combinations. Selected species-drug comparisons are shown in Supplementary Fig. S6.

Supplementary references

Clinical and Laboratory Standards Institute (CLSI), 2023. Breakpoint Implementation Toolkit: Accuracy Instructions for Prefilled Excel (Part E), version 1.0. https://clsi.org/media/i3ylyrde/part_e_accuracy_instructions_for_prefilled_excel_workbook.pdf (accessed 25 October 2026).

Clinical and Laboratory Standards Institute (CLSI), 2026. Performance Standards for Antimicrobial Susceptibility Testing. 36th ed. CLSI supplement M100. Clinical and Laboratory Standards Institute. https://clsi.org/shop/standards/m100/ (accessed 25 October 2026).

EUCAST, 2024. Reading guide for broth microdilution, version 5.0. https://www.eucast.org/fileadmin/eucast/pdf/MIC/Reading_guide_BMD_v_5.0_2024.pdf (accessed 30 September 2026).

EUCAST, 2026. EUCAST recommendations for MIC determination of aerobic and facultative anaerobic bacterial species with broth microdilution, version 1.0, September 2026. https://www.eucast.org/fileadmin/eucast/pdf/Document_Archive/bacteria/disk_diffusion_methodology/EUCAST_recommendations_for_broth_microdilution_MIC_determination_September_2026.pdf (accessed 21 October 2026).

Gitman, M., Chow, A., Hernandez, C., Mutnal, M., Henry, N., Teckenbrock, K., Chaudhary, S., Fey, P.D., Stevenson, J.S., Chan, Y.A., Pyden, A.D., Kearney, M., von Bredow, B., Ait Chebib, A., Garrett, E.M., Pu, M., Keelara, S., Remington, R., Carvalhaes, C., Lasky, C., Okeyo Owuor, T., Bobenchik, A.M., 2026. A comparative analysis of performance and time-to-result for VITEK REVEAL versus MicroScan during antimicrobial susceptibility testing for gram-negative bacteremia. Microbiology Spectrum 14, e01626-26. https://doi.org/10.1128/spectrum.01626-26

Jafari, M.J., Golabi, M., Ederth, T., 2024. Antimicrobial susceptibility testing using infrared attenuated total reflection (IR-ATR) spectroscopy to monitor metabolic activity. Spectrochimica Acta Part A: Molecular and Biomolecular Spectroscopy 304, 123384. https://doi.org/10.1016/j.saa.2023.123384

Loshchilov, I., Hutter, F., 2019. Decoupled Weight Decay Regularization. International Conference on Learning Representations. https://arxiv.org/abs/1711.05101

Riester, O., Kaiser, L., Laufer, S., Deigner, H.P., 2024. Rapid Phenotypic Antibiotics Susceptibility Analysis by a 3D Printed Prototype. Advanced Science 11, 2308806. https://doi.org/10.1002/advs.202308806

Rydzak, T., Groves, R.A., Zhang, R., Aburashed, R., Pushpker, R., Mapar, M., Lewis, I.A., 2022. Metabolic preference assay for rapid diagnosis of bloodstream infections. Nature Communications 13, 2332. https://doi.org/10.1038/s41467-022-30048-6

Sturm, A., Jóźwiak, G., Verge, M.P., Munch, L., Cathomen, G., Vocat, A., Luraschi-Eggemann, A., Orlando, C., Fromm, K., Delarze, E., Świątkowski, M., Wielgoszewski, G., Totu, R.M., García-Castillo, M., Delfino, A., Tagini, F., Kasas, S., Lass-Flörl, C., Gstir, R., Cantón, R., Greub, G., Cichocka, D., 2024. Accurate and rapid antibiotic susceptibility testing using a machine learning-assisted nanomotion technology platform. Nature Communications 15, 2037. https://doi.org/10.1038/s41467-024-46213-y

U.S. Food and Drug Administration (FDA), 2019. VITEK 2 AST - Gram Negative Imipenem: K183415. https://www.accessdata.fda.gov/cdrh_docs/pdf18/K183415.pdf (accessed 21 October 2026).

U.S. Food and Drug Administration (FDA), 2020. BioFire Blood Culture Identification 2 (BCID2) Panel: 510(k) substantial equivalence determination decision summary, K193519. https://www.accessdata.fda.gov/cdrh_docs/reviews/K193519.pdf (accessed 5 October 2026).

U.S. Food and Drug Administration (FDA), 2024. VITEK REVEAL GN AST Assay and VITEK REVEAL AST System: 510(k) substantial equivalence determination decision summary, K230675. https://www.accessdata.fda.gov/cdrh_docs/reviews/K230675.pdf (accessed 1 November 2026).
